# Nitrogen availability and plant functional composition modify biodiversity-multifunctionality relationships

**DOI:** 10.1101/2020.08.17.254086

**Authors:** Noémie A. Pichon, Seraina L. Cappelli, Santiago Soliveres, Tosca Mannall, Thu Zar Nwe, Norbert Hölzel, Valentin H. Klaus, Till Kleinebecker, Hugo Vincent, Eric Allan

## Abstract

The ability of an ecosystem to deliver multiple functions at high levels (multifunctionality) typically increases with biodiversity but there is substantial variation in the strength and direction of biodiversity effects, suggesting context-dependency. A better understanding of the drivers of this context dependency is essential to predict effects of global change on ecosystems. To determine how different factors modulate the effect of diversity on multifunctionality, we established a large grassland experiment with 216 communities, crossing a manipulation of plant species richness (1, 4, 8, 20 species) with manipulations of resources (nitrogen enrichment), plant functional composition (gradient in mean specific leaf area [SLA] to manipulate abundances of exploitative, fast-growing vs. conservative, slow-growing species), plant functional diversity (variance in SLA) and enemy abundance (foliar fungal pathogen removal). We measured ten above- and belowground functions, related to productivity, nutrient cycling and energy transfer between trophic levels, and calculated ecosystem multifunctionality. Plant species richness and functional diversity both increased multifunctionality, but their effects were context dependent. Species richness increased multifunctionality only when communities were assembled with fast growing (high SLA) species. This was because slow species were more redundant in their functional effects, whereas different fast species tended to promote different functions. Functional diversity also increased multifunctionality but this effect was dampened by nitrogen enrichment. However, unfertilised, functionally diverse communities still delivered more functions than low diversity, fertilised communities. Our study suggests that a shift towards fast-growing exploitative communities will not only alter ecosystem functioning but also the strength of biodiversity-functioning relationships, which highlights the potentially complex effects of global change on multifunctionality.

## Introduction

The ability of an ecosystem to deliver many functions at high levels simultaneously (i.e., multifunctionality) generally increases with diversity (Hector & Bagchi, 2007; Gamfeldt et al., 2008; Isbell et al., 2011; Lefcheck et al., 2015; Meyer et al., 2018; van der Plas, 2019). Although positive biodiversity-ecosystem multifunctionality relationships are common, studies have found substantial variation in the direction and strength of these relationships, which suggests that they depend on the biotic and abiotic environmental context (Balvanera et al., 2006; Soliveres, van der Plas, et al., 2016; van der Plas, 2019). However, only a few studies have attempted to understand this context dependency by determining how different drivers affect the biodiversity-multifunctionality relationship, and these have been either observational (Allan et al., 2015; Jing et al., 2015; van der Plas, 2019) or have focussed on only one of many potential drivers (Giling et al., 2019, but see Alsterberg et al., 2014; Liu et al., 2017). A better understanding of context dependency in biodiversity- multifunctionality relationships is important to predict effects of global change on ecosystems, because global change drivers, like nitrogen (N) enrichment or climate change, reduce biodiversity but also alter other aspects of the environment, and these changes may further alter the effects of biodiversity loss on ecosystem functioning. Experimental approaches are needed to rigorously test for context dependency but very few experiments cross diversity gradients with manipulations of major context drivers such as resources, natural enemies or plant species composition.

The main environmental factors influencing the strength and direction of the diversity- multifunctionality relationship are likely to be those that alter the mechanisms by which biodiversity affects functioning. Two of the main mechanisms behind the positive effects of plant diversity on ecosystem functioning are: i) complementary resource use amongst species (Fornara & Tilman, 2009; Jucker et al., 2015; Oram et al., 2018), and ii) increased enemy attack in species-poor communities, reviewed in Barry et al., 2019). If species vary in their resource uptake strategies, then diverse communities should use resources more efficiently (Tilman, 1982). Increasing resource levels would therefore remove opportunities for nutrient use complementarity between species and would equalise functioning between low and high diversity communities (Ratcliffe et al., 2017) (see Figure 1, Point 1). However, some experimental studies have shown neutral (Wacker et al., 2009; Hooper et al., 2012) or even positive effects of increased resource levels on the diversity-functioning relationship (Weigelt et al., 2009; Craven et al., 2016; Eisenhauer et al., 2018), suggesting that niche complementarity for soil nutrients does not always underlie diversity-functioning relationships. Nitrogen is one of the major soil nutrients limiting plant growth (Vitousek & Howarth, 1991) and differences in nitrogen use can be a key driver of niche differences between plants (McKane et al., 2002; Harpole & Tilman, 2005) and of positive effects of diversity on functioning (Fornara & Tilman, 2009). In addition, nitrogen enrichment is one of the major drivers of global change (Galloway et al., 2008; Battye et al., 2017) and testing how it modifies diversity effects is therefore of key importance to understand interactions between diversity loss and other aspects of global change. There is increasing evidence that natural enemy attack is also an important mechanism driving effects of diversity on function, and it can either enhance (Maron et al., 2011; Schnitzer et al., 2011), or reduce (Seabloom et al., 2017) the strength of the diversity-functioning relationship (Figure 1, Point 2). Foliar fungal pathogens, have received less attention than insects or vertebrate enemies but they are also key drivers of plant diversity and ecosystem functioning (Allan et al., 2010; Mordecai, 2011). There is little consensus on how plant enemies or resource availability affect diversity-functioning relationships but both could potentially be important drivers of context dependency.

**Figure 1:**
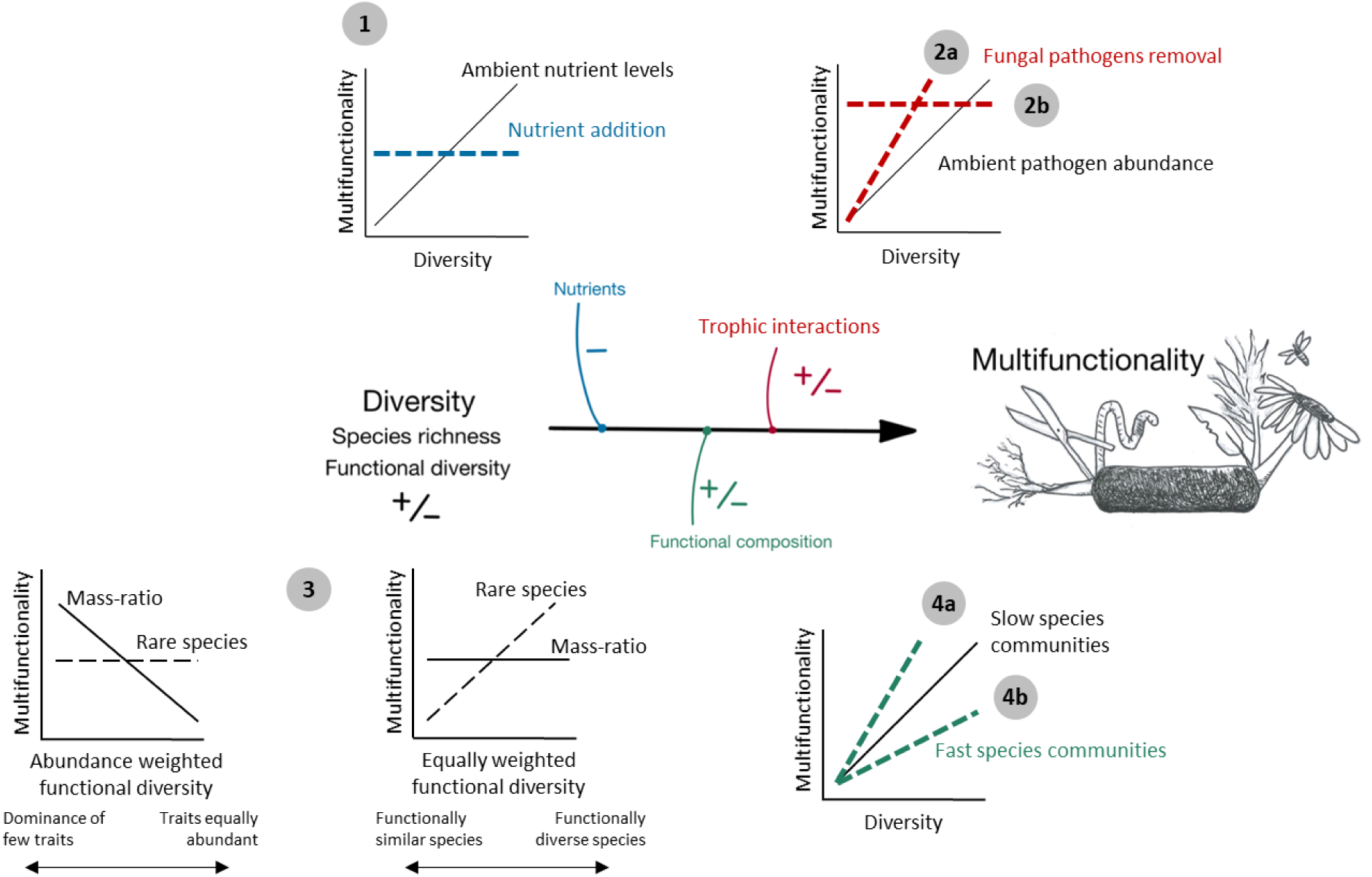
Concept of context dependency in plant communities. 1 Nutrient addition can reduce the potential for complementarity in diverse communities, and therefore dampen the diversity-multifunctionality relationship (Rattcliffe et al., 2017). 2a Pathogens can remove more biomass in diverse communities. Removing them would increase the diversity-productivity relationship (Seabloom et al., 2017). 2b Specialist pathogens preferentially attack low-diversity communities (Maron et al., 2011; Schnitzer et al., 2011). Removing them would dampen the diversity-productivity relationship due to higher functioning at low levels of diversity. 3 Functional diversity can be calculated in different ways. Weighting functional diversity by the species abundance assumes that a species contributes to functioning proportionately to its abundance, while equally weighted functional diversity assumes that the presence or absence of a species determines its effect on functioning. Mass-ratio effects, solid lines: if dominant species are key in supporting functioning, multifunctionality will decrease when abundance weighted functional diversity increases, and will not respond to equally weighted functional diversity. Importance of rare species, dashed lines: if rare species are increasing functioning, then multifunctionality will increase with equally weighted functional diversity, and will not respond to abundance weighted functional diversity. We show here the two extreme scenarios, but there are intermediate options between the importance of only mass-ratio or only rare species. 4a Fast species are more sensitive to resources and pathogen presence due to the growth-defence trade- off (Endara & Coley, 2011) and respond more strongly to environmental conditions: higher diversity- multifunctionality relationship can be more context-dependent in fast than in slow communities. 4b Slow species are more complementary in their resource uptake because they are adapted to low nutrient environments (Tilman & Wedin, 1991; Reich, 2014): lower diversity-multifunctionality relationship in fast communities than in slow.

In addition to the environment, the composition of the species pool (from which local plant communities are assembled) could modify biodiversity-multifunctionality relationships. A key element of composition could be the resource use strategies of the species, as those with different strategies might vary in their degree of resource use complementarity. The leaf economics spectrum (Wright et al., 2004; Díaz et al., 2016) distinguishes slow growing, conservative species from fast growing, acquisitive ones and is indicated by several traits, such as specific leaf area (SLA) and leaf nitrogen content. Slow species, with low SLA, are more competitive in the low nutrient environments they tend to dominate (Tilman & Wedin, 1991; Reich, 2014). They might therefore differ more in their competitive abilities for different nutrients, leading to greater nutrient use complementarity between slow species. In contrast, fast species, with high SLA, are abundant in high resource environments, where asymmetric light competition is prevalent (Hautier et al., 2009; Busch et al., 2019). Nutrient use complementarity between fast species might therefore be low. However, fast species are also likely to be more sensitive to enemies due to a growth defence trade-off (Endara & Coley, 2011), and enemies might therefore drive stronger diversity-functioning relationships in fast-dominated communities. Complementarity effects can lead to positive diversity-ecosystem functioning relationships for individual functions, which could also translate into higher multifunctionality in diverse communities. Such effects of the fast-slow functional composition of the species pool on diversity-function relationships would lead to different effects of diversity, depending on whether communities are dominated by fast or slow species, and therefore to interactions between changes in species composition and diversity (Figure 1, Point 4). Changes in diversity and fast-slow functional composition are typically correlated in observational studies (Allan et al., 2015) and new experimental approaches are therefore necessary to disentangle their interactive effects.

Biodiversity has many dimensions and different aspects of diversity can affect multifunctionality, with functional diversity being a key driver of functioning alongside species richness (Allan et al., 2015; Gross et al., 2017; van der Plas, 2019). A high fast-slow functional diversity could increase multifunctionality if species with different resource use strategies (fast vs slow) have complementary interactions (Cadotte et al., 2011; Handa et al., 2014). However, functional diversity can also reduce multifunctionality in harsh conditions where environmental filtering is strong and functionally distinct species are poorly adapted to the local conditions (Le Bagousse-Pinguet et al., 2019). In general, diversity and mass ratio effects can both occur and functions may in some cases be driven by complementary interactions between many less abundant species (Soliveres, Manning, et al., 2016), while in other cases they may be driven mostly by the dominant species, i.e. "mass ratio effects" (Grime, 1998) (Figure 1, Point 3). Functional diversity measures are strongly affected by rare species if only the presence/absence of species is considered but are mostly driven by differences between dominant species if they are abundance weighted. We therefore predict that non abundance weighted functional diversity should increase functioning when diversity effects and complementarity among rare species are important, e.g. because rare slow species provide novel functions (Mariotte, 2014). However, we would expect abundance weighted functional diversity to decrease functioning if mass ratio effects dominate and only the presence of key species is important (Mokany et al., 2008; Ruiz-Benito et al., 2014), e.g. because high productivity is high in communities dominated by fast species (Allan et al., 2015). Comparing effects of abundance vs. non abundance weighted fast-slow functional diversity might therefore shed light on the role of mass ratio vs. diversity effects in determining multifunctionality.

To better understand which contexts modify effects of diversity on multifunctionality, we measured multifunctionality in 216 plots of a grassland biodiversity-ecosystem functioning experiment. In this experiment, we tested the context-dependency of biodiversity-multifunctionality relationships to the addition of resources (N), enemy attack (fungal pathogens), and composition of species pool (functional composition). We factorially manipulated plant species richness, N enrichment, fungal pathogen removal (fungicide spraying), and plant fast-slow functional composition (by creating a gradient of abundance weighted specific leaf area [SLA]) and fast-slow functional diversity (the mean pairwise distance in SLA). We refer to these hereafter as functional composition and functional diversity. We therefore manipulated a specific element of functional composition and diversity in terms of the mean and variance in resource economics traits and did not explicitly manipulate composition and diversity in other trait axes. Our aim was to focus on the resource-economics axis because it has been identified as one of the key axes of plant functional variation (Wright et al., 2004; Díaz et al., 2016) and it represents a key set of traits predicting responses to N enrichment and effects on many ecosystem functions.

We measured ten functions on each plot, related to biotic interactions and biogeochemical cycling. We selected focussed on measures or proxies of process rates and consider high rates and low losses to indicate high functioning (Manning et al., 2018). We measured: above and belowground plant biomass, insect herbivore damage (hereafter herbivory) and foliar fungal pathogen infection, plant nitrogen and phosphorus uptake (quantified as ratios between available nitrogen and phosphorus content in the soil and nitrogen and phosphorus content in aboveground plant biomass), two enzymatic activities related to carbon (β-glucosidase) and phosphorus (phosphatase) cycling, soil respiration and soil carbon (C) storage, for more details see methods. We used a novel approach to analysing treatment effects on equal weight ecosystem function multifunctionality (Manning et al., 2018), in which we calculated multifunctionality at several thresholds between 50 and 80% of maximum function. We then combined all these multifunctionality measures in a single model (for details see methods), which allowed us to estimate the average effect of our treatments on multifunctionality, across thresholds. As any particular threshold is arbitrary, this allowed us to robustly quantify overall effects on multifunctionality. We hypothesised that: 1) species richness and fast-slow functional diversity will both increase multifunctionality, 2) the strength of the diversity effect will depend on the functional traits of the species which are combined (i.e. the functional composition of the species pool), and 3) increased resource levels and removal of fungal pathogens will decrease the effect of diversity on multifunctionality.

We also included in the analyses possible interactions, not represented here, between the different factors (nutrient and pathogens for instance) and with more than two factors.

## Results

### Richness-multifunctionality relationships depend on functional composition

Diversity effects on multifunctionality were highly context-dependent in our experiment, as we found three interactions between effects of richness and other factors. The strongest of these was the interaction between SLA and richness (estimate of the interaction: 0.025 +/- 0.009 SE), which meant that the effect of sown species richness on multifunctionality was modified by fast-slow functional composition. These results were consistent when using multiple trait measures of functional composition instead of just SLA (Table S1). Species richness decreased multifunctionality slightly in communities with low mean SLA but increased multifunctionality in high mean SLA communities (Figure 2a and b and Table S2). Species richness therefore had a positive effect on multifunctionality in communities assembled from fast-growing, acquisitive species, but a neutral to negative effect when the species pool contained only slow-growing, conservative species. At the same time, SLA generally increased multifunctionality (and the main effect of SLA was the single strongest driver of multifunctionality) but the interaction with richness meant that SLA increased multifunctionality more strongly in diverse communities. Subsequent analyses revealed that these results were not driven by any particular function (Table S3 and Figure S1) and remained consistent regardless of the number of functions included in our multifunctionality metric (Figure S2 and Figure S3), or the use of the realised [as opposed to the sown] species richness (Table S4). SLA as a main effect had contrasting effects on individual functions, with a positive effect on functions measured aboveground (pathogen presence, herbivory), and a negative effect belowground (belowground biomass, phosphatase activity, soil respiration, Figure 3). This shows that fast and slow species drive different functions. We further explored whether the different functional effects of fast vs. slow species may explain why diversity has different effects on multifunctionality in slow vs. fast communities (see below).

**Figure 2:**
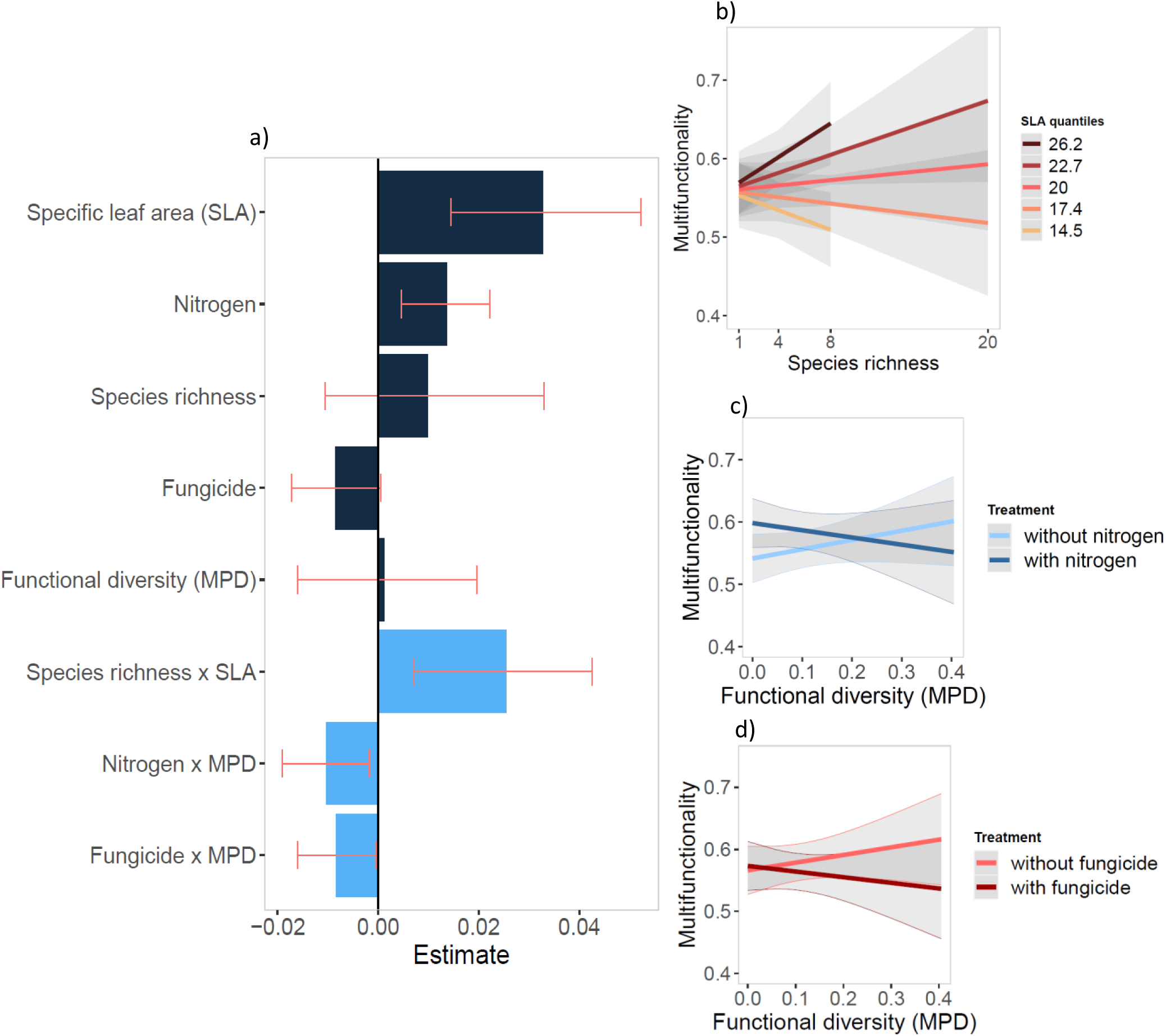
Context-dependent diversity-multifunctionality relationships. a) Standardised effect (+/- confidence intervals) of each scaled centred variable on multifunctionality, across seven thresholds. Only significant effects are shown to facilitate interpretation. All main effects are shown regardless of significance, as they are all part of at least one significant interaction. b) Interaction between sown species richness and community weighted mean specific leaf area (SLA, m^2^ kg^-1^), represented as quantiles of SLA (see raw data in Figure S4). c) Interaction between functional diversity (MPD, mean pairwise distance) and nitrogen enrichment. d) Interaction between functional diversity and fungicide application. N = 216 plots. Details of the model output can be found in Table S2. Species richness is here sown richness.

**Figure 3:**
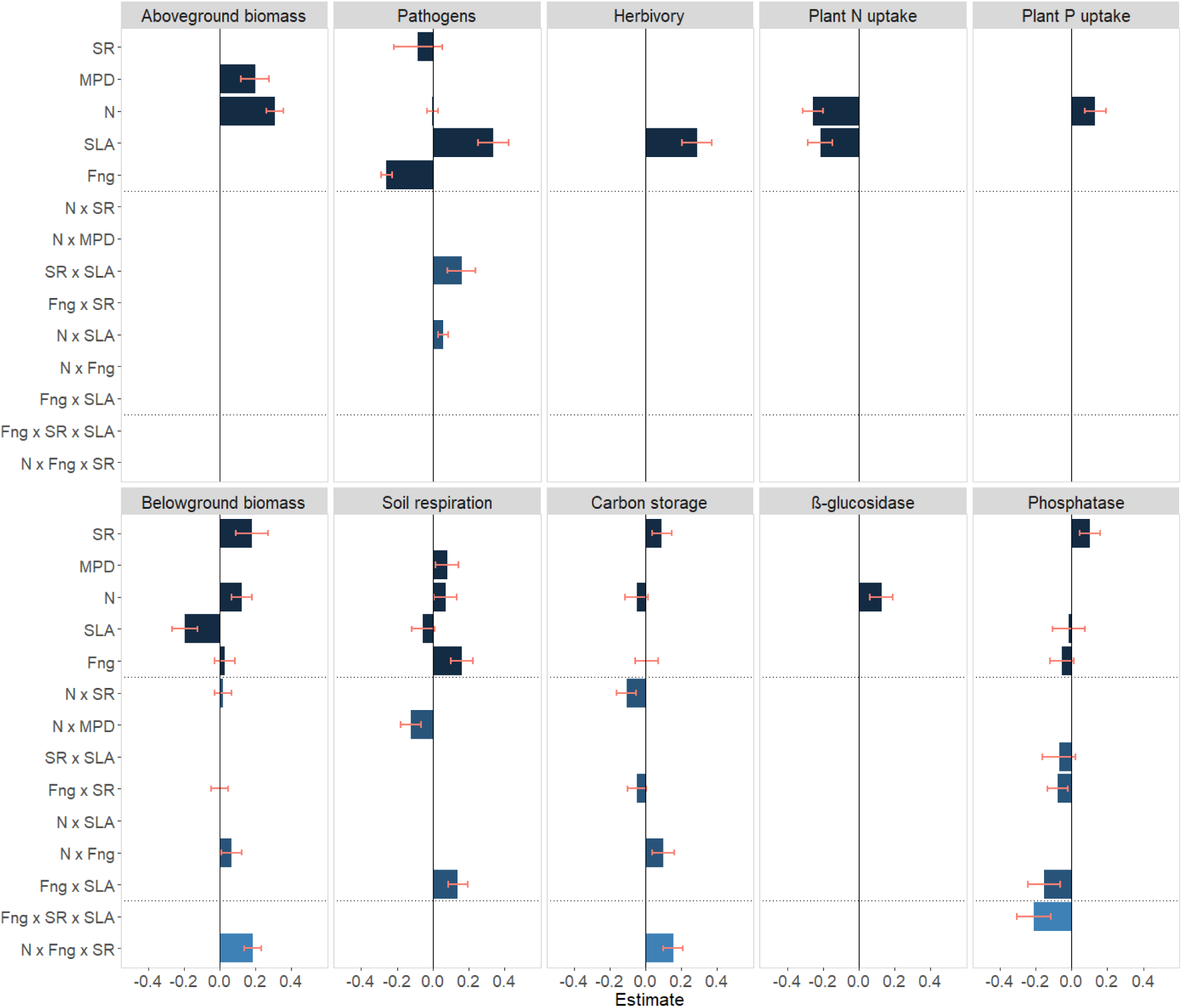
Drivers of individual functions. Mean standardised effect (+/- confidence intervals) of each scaled and centred variable on individual functions after model simplification. Only significant effects are shown but main effects and interactions which are part of higher order interactions are displayed regardless of significance, see Figure S1 for the full model. SR: sown plant species richness, MPD: mean pairwise distance in SLA equally weighted, N: nitrogen enrichment, SLA: abundance weighted community specific leaf area, Fng: fungicide treatment. Each function was measured on 216 plots. See Table S9 and Figure S1 for details.

### High resource levels dampen effects of functional diversity on multifunctionality

Sown fast-slow functional diversity, calculated as the mean pairwise distance between species in their SLA, increased multifunctionality in unfertilised plots (Figure 2a and c). However, nitrogen enrichment dampened this positive effect because it increased multifunctionality in plots with low functional diversity (estimate of the interaction: -0.010 +/-0.004 SE). Nitrogen enrichment increased soil respiration, in particular, in plots with low functional diversity (Figure 3). N enrichment also increased several individual functions related to plant productivity and soil microbial activity overall (above and belowground biomass, soil β-glucosidase activity, plant P uptake, see Figure 3). Interestingly, N enrichment also interacted with species richness to affect belowground biomass and C storage (Figure 3), but this interaction further depended foliar pathogen removal (fungicide treatment).

Foliar pathogen removal also dampened effects of fast-slow functional diversity on multifunctionality, as functional diversity only increased multifunctionality where foliar pathogens were not reduced by fungicide application (estimate of the interaction: -0.008 +/-0.004 SE, Figure 2a and d). However, this interaction should be treated with caution as it seems to be driven mainly by the response of fungal pathogen damage (see Table S5), which was reduced by fungicide, as expected. Fungicide also increased total soil respiration in high SLA communities and belowground biomass in species-rich, fertilised communities and was also involved in some other complex interactions for C storage and phosphatase.

### Effect of change in species abundances on functional diversity

Although all species started at equal abundance in our experiment, species shifted their relative abundances over time. To test the effect of these abundance changes on functional diversity, we calculated abundance weighted mean pairwise distance (MPDw), weighted by realised species abundances. While sown functional diversity increased multifunctionality, the abundance-weighted functional diversity showed the opposite pattern (Figure 4). These contrasting effects indicate that the communities with the highest multifunctionality were those which had been sown with functionally distinct species (i.e. with high unweighted MPD), but in which functional diversity decreased because they became dominated by species with similar functional trait values (they ended up with lower abundance weighted MPDw). Such high multifunctionality communities are dominated by functionally similar species but contain functionally distinct rare species.

**Figure 4:**
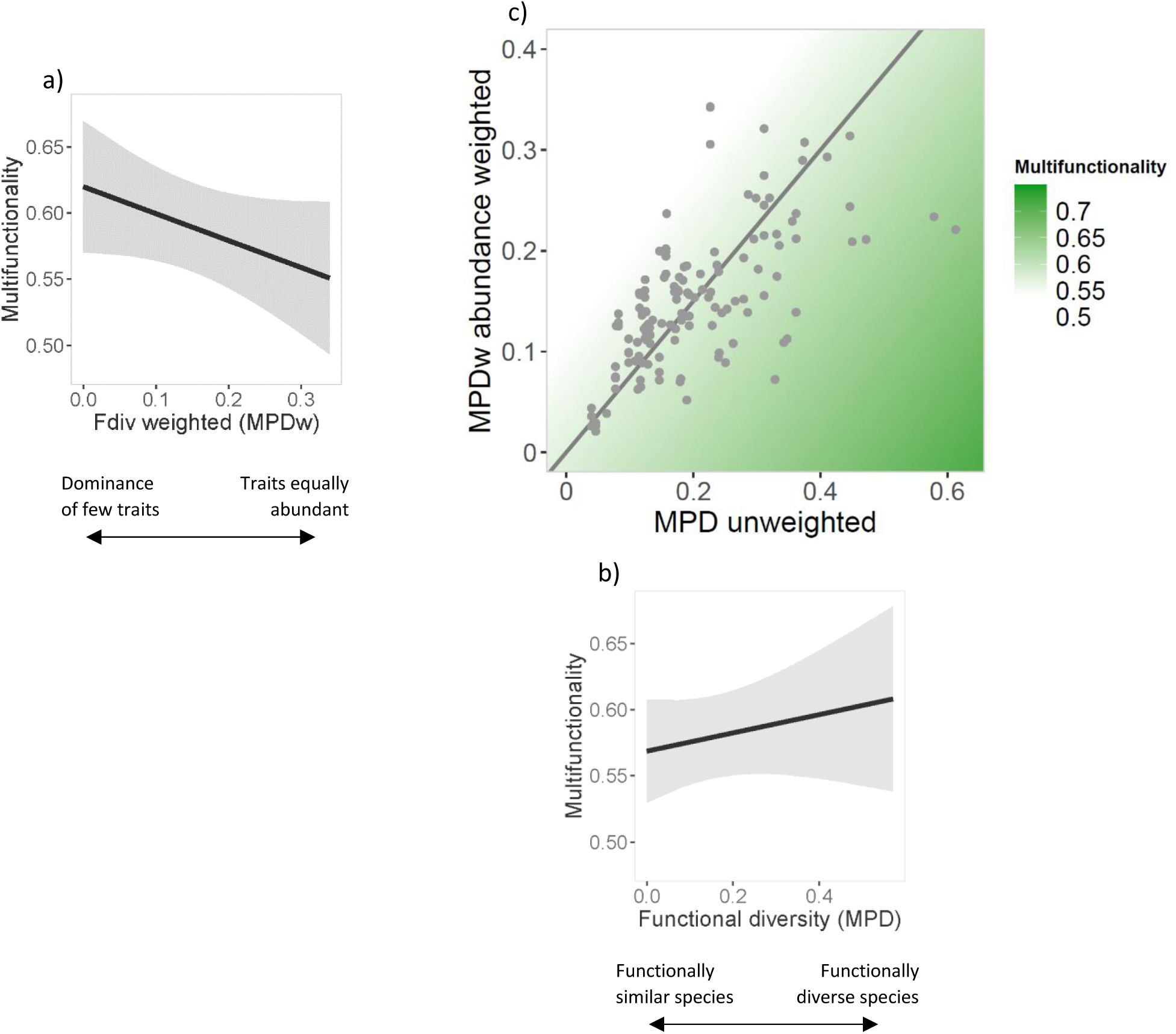
Effect of sown vs. realised functional diversity on multifunctionality. Effect (+/- confidence intervals) on multifunctionality of a) abundance weighted, realised, functional diversity (MPDw) and b) non-abundance weighted, sown, functional diversity (MPD). c) MPDw plotted against MPD. The green area represents the model-predicted multifunctionality from a 65% threshold (midpoint between 50% and 80% thresholds), but the effect is similar at other thresholds, as there is no interaction between threshold and MPD. The high multifunctionality plots are those with both a relatively high MPD and a relatively low MPDw (bottom right corner). Left of the grey line: functional diversity increases from sown communities. Right of the grey line: functional diversity decreases from sown communities. Monocultures were included in the individual models but are not shown in c) as they are all coded as 0 MPD and MPDw. See Table S10 for details on the model output.

### Effects of individual species on functions

The positive effects of both SLA and functional diversity on multifunctionality show that combining multiple fast-growing species in a community leads to high multifunctionality, whereas combining slow species does not increase multifunctionality. There are two main factors that could cause this. First, if slow species were more redundant in their functional effects (they all drive similar functions), then adding more slow species to a community (increasing diversity of slow species) would not increase multifunctionality. Second, slow species could have trade-offs in their functional effects (i.e. they increase some functions and reduce others), while fast species have more synergies (positive effects on multiple functions). This would mean that increasing diversity of slow species would not enhance multifunctionality in the temperate grassland we studied, because adding a slow species to a slow community would tend to reduce as many functions as it enhanced.

To test these ideas, we calculated the effect of our 20 species on each function, by fitting regression models for each function with the abundance of each species as predictors (following the approach used in Hector & Bagchi, 2007; Isbell et al., 2011). We extracted the significant positive and negative effects of each species on each function. To explore how combining sets of slow or fast species would affect multifunctionality, we calculated the dissimilarity in functional effects between sets of four species (see methods for more details). This showed that fast species were more dissimilar in the functions they drove (Table S6, Table S7 and Table S8), as an increase in mean SLA between species increased the dissimilarity in the set of functions they affected. The results were similar regardless of the use of presence-based or abundance weighted effects, the number of species considered when calculating dissimilarity, and for both positive and negative effects (see methods and Tables S5 to S7), showing that all slow species tend to affect similar functions, while fast species affect different functions. For instance, slow species like *Achillea millefolium, Centaurea jacea, Festuca rubra* and *Salvia pratensis* all drove biomass production and insect herbivory, while fast species tended to drive different functions, e.g. *Crepis biennis* increased N uptake, while *Holcus lanatus* increased pathogen infection.

In order to test whether there are also more trade-offs in functional effects between slow species, we calculated, for each pair of species, the proportion of functional effects that were trade-offs (see methods). The effects were non-significant, both when analysed based on species presence or on species abundance (Table S6). It therefore seems that trade-offs in functional effects are not more common for slow species and that instead slow species are more redundant in their functional effects because they tend to drive similar functions.

## Discussion

### Functional trait composition modulates richness effects on ecosystem functioning

Our results show that there is substantial context dependency in biodiversity-multifunctionality relationships. The most important driver of this context-dependency in our study was the functional composition of the species pool, in terms of whether communities were composed of fast-growing acquisitive species or slow-growing conservative species. When experimental (Hector et al., 2011; Craven et al., 2018) or observational (e.g. Allan et al., 2015) studies have considered compositional and diversity effects together, they have compared their relative importance but not potential interactions between them. By experimentally crossing manipulations of fast-slow functional composition and of species richness, we were able to show that community SLA and species richness had an interactive effect on multifunctionality, which means that composition and diversity are not always independent drivers of multifunctionality. Some studies have considered how the functional *diversity* of the species pool impacts species richness-functioning relationships (Ratcliffe et al., 2017; Wagg et al., 2017) but our results show that the functional *composition* of the species pool may be a stronger driver. Two different analyses provided evidence that multifunctionality was affected by both mass ratio, or selection, effects and diversity, or complementarity, effects and that these interacted. The analysis of individual species effects on functions showed that certain slow species drove many functions and at the same time abundance weighted functional diversity reduced multifunctionality, indicating mass ratio effects because multifunctionality was highest when the dominants were functionally similar. In contrast, fast species drove different functions, suggesting complementarity between them, and non-abundance weighted functional diversity increased multifunctionality, indicating that functionally distinct rare species also provided different functions. Our results therefore highlight that while species richness and composition have usually been seen as alternative or opposing drivers of functioning (i.e. mass-ratio vs. diversity effects, Tilman, 1997; Grime, 1998; Mokany et al., 2008; Ruiz-Benito et al., 2014), they can modify each other’s effects (Craven et al., 2018; Le Bagousse-Pinguet et al., 2019).

Our analysis of individual species effects on functions showed that slow species were more redundant in their functional effects, because some slow species (especially *Achillea millefolium, Centaurea jacea, Festuca rubra* and *Salvia pratensis*) drove multiple functions, while fast species tended to affect a different function each. Therefore, combining multiple fast species increased multifunctionality because each species increased a different function, while combining multiple slow species did not add more functions, as they were more redundant in the functions they provided. We also found that fast and slow species were complementary with each other and provided different functions, also see below. At the same time, some slow species had only negative effects on functioning, meaning that on average monocultures of slow species had low multifunctionality. This suggests that amongst slow species there might be a “multifunctional selection effect” with high multifunctionality driven by the abundance of those species that promote several functions, while amongst fast species there is “multifunctional complementarity” with different species driving different functions. This idea is further supported by the fact that diversity x composition interactions were not significant for most individual functions, as opposed to what we found for multifunctionality. Instead, species richness increased belowground biomass and carbon storage regardless of community SLA, and phosphatase activity peaked in plots dominated by slow species, regardless of their richness. Diversity therefore did increase several individual functions within slow species communities, presumably because the slow species were complementary in their effects on these individual functions (Cappelli et al., 2022), but this complementarity was not realised when combining multiple functions.

Here, we focussed on the fast-slow resource use axis as a key element of functional composition (which was well represented by the usage of SLA as a unique trait; Table S1). However, other functional traits or axes, such as size (Díaz et al., 2016), dispersal and root strategies (Bergmann et al., 2020) also determine overall functional composition. Future work manipulating these elements of functional composition in combination with diversity would be very valuable.

### Resource levels modulate functional diversity effects on ecosystem functioning

In addition to sown species richness, sown functional diversity was an important driver of multifunctionality but its positive effect was dampened by N enrichment. This results is at odds with reports of stronger (Weigelt et al., 2009) or consistent (Craven et al., 2016) diversity-productivity relationship in response to increased resource supply. There are three main reasons why our results may differ from these previous findings. Firstly, most other studies added multiple nutrients, which increased total resource levels, whereas we added nitrogen only which may have led to an unbalanced nutrient stoichiometry and therefore a reduction in complementarity (Cardinale et al., 2009). Secondly, N addition may have a larger effect on the diversity-*multifunctionality* relationship than on the diversity-*productivity* relationships examined in previous studies. Consistent with this, our study also found that the effect of diversity on aboveground productivity was independent of fertilisation (Figure 3), and that the interaction between N and either species richness or functional diversity was inconsistent across individual functions. Further, a study on soil multifunctionality (Eisenhauer et al., 2018), also showed that the positive diversity effect on multifunctionality was weaker under N enrichment. Finally, we only found that N modified the effect of functional diversity, not of species richness, and it may be that N addition is more likely to reduce effects of fast-slow functional diversity than of species richness because the fast slow continuum is more closely linked to resource use. Overall, our results show that N addition only increases functioning in low diversity communities while adding N to functionally diverse communities would not further increase multifunctionality. However, high diversity, unfertilised communities still delivered higher multifunctionality than low diversity, fertilised communities (0.624 and 0.588 respectively), suggesting low input, high diversity grasslands could deliver high overall multi-functioning relative to intensive systems.

Foliar fungal pathogen removal also reduced diversity-multifunctionality relationships and modified the diversity effect on individual functions. Previous studies have shown that specialist soil pathogens can drive diversity-productivity relationships by reducing biomass particularly at low diversity levels (Maron et al., 2011; Schnitzer et al., 2011). In our experiment, reducing foliar fungal pathogen abundance reduced multifunctionality in functionally diverse communities, suggesting that fungal endophytes present in diverse plant communities may boost certain functions. Fungicide application might alter the composition of pathogen communities by removing specialists (Cappelli et al., 2020), but it might also reduce the abundance of mutualists (fungal endophytes, Karlsson et al., 2014; Ayesha et al., 2021), particularly in diverse plant communities. A high diversity of bacterial leaf endophytes has been shown to drive effects of tree diversity on ecosystem functioning (Laforest-Lapointe et al., 2017), and fungal endophyte diversity might therefore also be important. However, the interaction between fungicide and diversity was sensitive to the inclusion of pathogen damage in the multifunctionality measure, even though functional diversity and fungicide application did not interact to affect pathogen damage itself, meaning the effect of pathogen removal on the diversity-multifunctionality relationship should be treated with caution. Nevertheless, our results indicate that fungal endophyte communities may deserve more attention as potential drivers of diversity-multifunctionality relationships.

### Both dominant and rare species increase multifunctionality

Multifunctionality was maximised at high levels of sown functional diversity but was reduced by high levels of abundance weighted functional diversity. These contrasting effects of non-abundance and abundance weighted functional diversity mean that multifunctionality was highest when communities became dominated by one or a few functionally similar species, but still retained some distinct rare species (relatively low abundance weighted functional diversity, combined with high sown functional diversity; Fig. 4). The negative effects of abundance weighted functional diversity mean that multifunctionality was maximised when the dominants in a community were functionally similar, which suggests that mass-ratio effects were important for increasing multifunctionality, this is analogous to a multifunctional selection effect where species with certain traits are key for providing multifunctionality. Studies in biocrust systems (Maestre et al., 2012) and global drylands (Le Bagousse-Pinguet et al., 2019) have also shown that multifunctionality was maximised when communities were dominated by functionally similar species, which were well adapted to the environmental conditions. In our experiment, communities dominated by high SLA species could enhance multifunctionality (CWM SLA had the strongest effects on multifunctionality in our study) by enhancing functions related to fast biogeochemical cycling (herbivory, pathogen infection, see Figure 3). At the same time some individual slow species were also able to provide many functions. On the other hand, the increase in multifunctionality with non-abundance weighted functional diversity emphasises the importance of diversity and functionally unique species in driving multifunctionality (McLaren & Turkington, 2010; Vogel et al., 2019). Our analysis of individual species effects on multifunctionality indicated that fast species tended to be more complementary in the functions they drove, but this result indicates that fast and slow species are also complementary and provide different functions from each other. These findings are also in accordance with a body of literature on the importance of locally rare species for the provision of multiple ecosystem functions (Lyons et al., 2005; Mouillot et al., 2013; Soliveres, Manning, et al., 2016). In our experiment the presence of low SLA species led to an increase in belowground biomass (Figure 3) and soil respiration (on plots without fungicide), perhaps indicating increased microbial activity in slow growing communities (Delgado-Baquerizo et al., 2018). The addition of a few rare slow species could therefore have increased the levels of certain soil functions and could have boosted multifunctionality in otherwise fast dominated communities (Le Bagousse-Pinguet et al., 2021).

### Conclusions

Our results shed light on the context dependence of diversity effects on multifunctionality. The most important driver of context dependency in our study was the functional trait composition of the species pool (whether communities were assembled with fast or slow species). Our findings show that shifts in fast-slow functional composition due to global change will not just alter ecosystem functions themselves but will also alter the strength of diversity-functioning relationships. This could result in complex interactions between global change drivers, for instance, if some drivers alter fast-slow functional composition and others change diversity. In addition, they suggest that managing functional composition can buffer the impacts of other global change drivers on ecosystem functioning and biodiversity-functioning relationships. For the functions considered here, the optimal community for multifunctionality contained several different fast species at high abundance and some functionally distinct slow species at low abundance. However, the optimal community is likely to differ for other sets of functions or environmental conditions, and considering which functions should be maximised will be important to define optimal functional composition and diversity for a given scenario. Overall, these findings underline the need to consider multiple diversity dimensions, and interactions between them and abiotic conditions, to optimise multifunctionality in the context of multiple simultaneous global change factors.

## Material and methods

### The PaNDiv Experiment

We set up a field experiment in October 2015 in Münchenbuchsee (Switzerland, 47°03′N, 7°46′E, 564 m a.s.l.) with factorial manipulations of plant species richness, plant functional composition, N enrichment and foliar fungal pathogens (Pichon et al., 2020). The site had been unfertilised for at least 10 years before the start of the experiment. The soil is characterized as 0.7 to 1 m deep brown earth (Cambisol), according to the Geoportal of the Canton Bern (http://www.geo.apps.be.ch). We measured total soil N and C concentrations and pH in the top 20 cm of soil at the start of the experiment and found concentrations of 2.3-4.2% C, 0.26-0.43% N and a pH of 7.4. The original grassland was dominated by *Dactylis glomerata, Lolium multiflorum, Crepis biennis, Plantago lanceolata* and *Trifolium sp*. The area was cleared from vegetation and ploughed before the establishment of experimental plots.

The whole experiment consisted of 216 plots of 2 m × 2 m separated from one another by a 1 m path and arranged in four blocks. We sowed the different communities with enough seeds to obtain a density of 1000 seedlings per m^2^, equally divided between the species and corrected for species germination rates. The species pool consisted of 20 herbs and grasses typical of mesic, Central European grasslands (see list of species Table S11). We divided the species into two pools corresponding to different resource economic strategies (Wright et al., 2004), using literature values of leaf N content and specific leaf area (SLA) to classify the species. We created a pool of 10 species with high SLA and leaf N, which have a fast-growing, acquisitive resource use strategy, with short leaf lifespans and high photosynthetic rates, and a pool of 10 species with a more slow-growing, conservative strategy, long leaf lifespans and low photosynthetic rates. For simplicity we refer to these species as “fast” and “slow” throughout. We excluded legumes from the species pool as they are expected to decline with N enrichment and including them in the slow growing pool only, would add an additional difference between the pools. We therefore might have underestimated the diversity effects by excluding legumes from the species pool.

All 20 monocultures were established, together with 15 four species, 15 eight species and 4 twenty species plots. Plots with four and eight species could contain either only fast or only slow species, or a mix of the two strategies. This allowed us to create a large gradient in functional composition (mean SLA) and functional diversity (variance in SLA), independent of species richness. Species compositions for each plot were randomly selected from the respective species pool, with the constraint that all plots (except monocultures) had to contain both grasses and herbs. Each particular species composition (54 in total) received the four combinations of N enrichment and fungicide application, for a total of 216 plots. Fertilised plots received N in the form of urea twice a year in April and late June, for an annual addition of 100 kg N ha^-1^y^-1^, which corresponds to intermediately intensive grassland management (Blüthgen et al., 2012). Urea addition had no significant effect on soil pH measured in autumn 2017 (Table S12). In order to manipulate multitrophic interactions we removed foliar fungal pathogens using fungicide. The plots were sprayed with two fungicides to try to achieve broad suppression of the fungal community ("Score Profi", 24.8 % Difenoconazol 250 g.L^-1^ and "Ortiva", 32.8% Chlorothalonil 400 g.L^-1^ 6.56% Azoxystrobin 80 g.L^-1^), four times during the growing season (beginning of April and June, late July and September). The same amount of water was applied to the untreated plots at the same time. The fungicides were very effective in suppressing rusts and mildews but less effective in reducing infection by leaf spots, and had very few non target effects (Cappelli et al., 2020). The experiment was weeded three times a year to maintain the diversity levels. Realised species richness values were very close to sown values (Pearson’s correlation = 0.997). All plots were mown twice a year, in mid-June and mid-August, which corresponds to intermediate, to extensive, grassland management in the region.

Further details about the site characteristics and experimental setup of the PaNDiv Experiment can be found under Pichon et al. (2020).

### Measurements on the plant community

#### Plant traits

We measured specific leaf area (SLA = surface/dry weight, m^2^ kg^-1^) in August 2017 and in June and August 2018, by taking one leaf from each of five individuals in all 80 monoculture plots. We sampled treatment-specific monocultures to include effects of nitrogen addition and pathogen removal on trait expression. We rehydrated the leaves and measured leaf surface, and dry weight according to the protocol of Garnier et al. (2001). The SLA per species is the average of the trait values across the five samples and across the three sampling dates and therefore averages across seasonal variation. We used only SLA to calculate functional diversity and composition as our experimental design involved manipulating the mean and variation in SLA within communities (species were defined as fast and slow based on SLA). However, we also tested how well SLA represented the overall resource economics spectrum by using a combined strategy index based on SLA, LDMC and nitrogen concentration in plant aboveground biomass, see Table S1.

#### Functional composition (CWM SLA)

We visually estimated the cover of each plant species (target species and all weeds together) before mowing. We calculated the community weighted mean (CWM) SLA per plot using the relative cover estimates averaged across 2017 and 2018: CWM = ∑ *pi* * *xi*; with *pi* the relative abundance of the species *i* and *xi* the trait value of *i* in each monoculture of the same treatment (i.e. nitrogen and fungicide added or not).

#### Functional diversity (MPD and MPDw)

As a measure of functional diversity, we calculated a community mean pairwise distance, as the mean distance in SLA between the species sown in the plot (de Bello et al., 2016). This represents the initial (sown) functional diversity of the plots when species were at equal abundance. The SLA values used were the monoculture traits of the same treatment, and all monocultures had an MPD value of 0 by definition.

To assess how changes in species relative abundances affected the realised MPD of the experimental communities, we calculated an abundance weighted index, MPDw, using cover values from 2017 and 2018 as abundance measures. The two indices were highly correlated (see Figure S5), partly due to coding the monocultures as 0. To avoid issues of multicollinearity, which could bias regression coefficients, we therefore analysed the effect of MPD and MPDw in two separate models (Table S10). Model output calculated with non-abundance weighted MPD, but excluding the species that were not recorded at least once between 2017 and 2018 did not differ from the main model output (see Table S4).

### Ecosystem functions

#### Selection of functions

We measured ten functions related to resource and energy transfer both above and belowground: above and belowground biomass production, foliar fungal pathogen infection, insect herbivore damage, N and P uptake, soil respiration, C storage, and beta-glucosidase and phosphatase activity. Half the functions were characteristic of aboveground processes (aboveground biomass, pathogens, herbivores, N and P uptake), and half of belowground processes (belowground biomass, soil respiration, C storage, beta-glucosidase, phosphatase). The functions were related to primary production (above and belowground biomass), to secondary production (pathogens, herbivores), and to microbial activity (soil respiration, beta-glucosidase, phosphatase). The functions included measures of nutrient cycling (N and P uptake, phosphatase) and C cycling (soil respiration, C storage, beta-glucosidase). In addition, we aimed to measure fluxes not stocks, as we were interested in the transfer rate within the ecosystem (Barnes et al., 2018) and we consider high rates of transfer and low loss rates to indicate high functioning. We therefore calculated N and P uptake instead of soil available N and P stocks. Two functions could be considered as stocks, i.e. belowground biomass and C storage, but they can also be seen as fluxes over several years, as they measure the difference in C stock and in belowground biomass between the end and the start of the experiment. Pathogen and herbivore damage are measures of energy transfer to higher trophic levels and high levels of these therefore indicate high functioning because they mean that energy is transferred more rapidly between trophic levels and that secondary production is higher (Meyer et al., 2015; Barnes et al., 2018; Manning et al., 2018; Wang et al., 2019; Barnes et al., 2020; Buzhdygan et al., 2020). Soil respiration can be considered either as an indicator of high functioning (increased soil activity) or as an indicator of low functioning (C loss from the system). Due to the positive correlation between respiration and plant biomass (Figure S8), we chose to consider soil respiration as indicating high functioning. However, the results were not altered when we reversed respiration values to consider high respiration to be low functioning, see Table S14. Our results are therefore robust to the uncertainty in how to consider soil respiration.

We also insured that all functions were not strongly correlated with each other and that no one function strongly correlated with multifunctionality (Fig. S6). The functions described in this study were all measured between June 2017 and autumn 2018. For each function measured more than once, we used the mean value per plot over the whole period (see Table S9).

#### Aboveground biomass

We collected aboveground biomass twice per year, before the mowing. The samples were taken by clipping plant material to 5cm above ground level, in two quadrats of 20 cm × 50 cm located in the centre of each plot. We weighed the biomass after 2 days of drying at 65°C. The plot target biomass production (i.e. of the sown species without weeds) was calculated by multiplying the percentage cover of weeds by the total biomass and subtracting this estimated weed biomass from the total weight. To determine if the estimation was accurate, we sorted the biomass per species in June 2017 for all plots of one block. The sorted target biomass (without the weeds) was highly correlated with the estimated target biomass (R^2^ = 0.994, Figure S7). The mean weed percentage was around five percent and therefore removing the weeds made little difference to the total biomass (Pichon et al., 2020).

#### Herbivory

Herbivory was assessed in May and August 2018. Five individuals of each target species were randomly selected from each plot and five leaves per individual were assessed for damage. Leaves were selected from the middle tier of each individual, and juvenile and senescent leaves were excluded. Damage was characterised by the type of herbivory; chewing, sucking, leaf mining and rasping damage based on the methods of Loranger et al. (2014). In this study, we calculated the community weighted mean total damage using the sum of damage types per leaf, per species, weighted by the cover of the particular species.

#### Pathogens

Fungal infection was measured per species per plot once in October 2017 and twice in 2018, in July and October. Ten randomly chosen individuals in the central square meter of the plots were visually screened for signs of infection and the percentage of infected individuals was recorded. If there were less than 10 individuals in the central square meter, individuals in the rest of the plot were screened. If there were fewer individuals, the percentage of infected individuals was calculated based on all individuals present. The species level infection and the percentage cover data were used to calculate a community weighted mean of fungal infection per plot and season.

#### Plant N and P uptake

We calculated plant N and P uptake using total biomass and soil available N and P content. The biomass collected in June and August 2018 was homogenised per plot and we ground a minimum of 5g per plot with a cyclone mill to obtain a fine powder. We then used near infrared reflectance spectroscopy (NIRS) analyses to estimate biomass N and P content using calibrations by Kleinebecker et al. (2011). Soil samples were collected five times in 2018 using two homogenised cores per plot. Soil NO ^-^, NH_4_ ^+^ and PO_4_ ^3-^ were extracted with CaCl and analysed using a Continuous Flow Optical- Absorption Spectrometer (CF-OAP; model Skalar Scan+; Skalar Analytical, Breda, The Netherlands). The plant N and P uptake are the mean plant N and P content divided by the mean available N (NO ^-^ plus NH ^+^) and available P (PO_4_ ^3-^), respectively.

#### Enzymatic activity

The activity of two enzymes related to carbon and phosphorus cycling, β-glucosidase and phosphatase, was measured in autumn 2017 and in April, May, July and August 2018. These functions give an estimate of soil microbial activity, and β-glucosidase has been shown to be a good soil health indicator (Turner et al., 2002). We took two soil samples per plot of 1.5 cm diameter to 20 cm depth. The fresh samples were homogenised and sieved through 2mm meshes. In each sample, we measured the β-glucosidase and phosphatase enzymatic activities, following the protocol of Tabatabai (1983). For the analyses presented here, we calculated the mean value of each enzymatic activity for all the periods sampled.

#### Belowground biomass

Root biomass was measured once in autumn 2017. We took two cores per plot to 20 cm depth (440 cm^3^ of soil). We homogenised the two samples and used a subset of 40 g fresh soil in which we sorted out the roots. The samples were washed in 200 µm sieves and the roots sorted out with tweezers. We dried the roots at 65°C for 48 hours. We estimated soil bulk density by weighing 40g of soil from the same plot before and after drying for 24 hours at 105°C. Bulk density did not differ between the treatments (Table S13). The final belowground biomass per plot is the weight of roots per g of dry soil.

#### Respiration

We measured overall soil respiration five times during 2018. We placed one PVC ring per plot at least a week before taking the measurements. The vegetation was clipped before starting the measures without disturbing the soil. We took two measures per plot and repeated the measures in case of large difference between the respiration rates. We took the measurements using a soil CO2 flux chamber (6400-09) attached to a LI-6400XT (Li-Cor Inc.). Soil respiration varies during the day and between days. To minimise this variation, we took the measurements starting at 10 am and finishing the latest at 2 pm, as this time span is expected to provide rates representative of the mean daily respiration (Mielnick & Dugas, 2000; Castillo-Monroy et al., 2011). We measured one block per day (54 plots), avoided humid days and kept the measurement days as close to each other as possible (max 6 days for one sampling session). The final soil respiration measure is the mean measure per plot.

#### Carbon storage

We measured soil carbon concentration on soil samples of 440 cm^3^, taken to 25 cm depth in autumn 2017. We took two samples per plot, homogenised them and removed stones and living material (roots, fauna). We weighed the samples before and after drying at 65°C for 48 hours. To calculate C storage, we compared the 2017 samples to soil samples taken in autumn 2015 before the start of the experiment. In 2015, we took 5 soil samples per plot from a subset of 89 plots distributed across the field. As the field had just been ploughed and homogenised there was relatively little variation between adjacent plots and we estimate soil C concentrations for all plots using kriging (package geoR, estimating sill and range of the data variogram using the variofit function). Analyses of soil total C were conducted in a CNS Analyser at the Institute of Geography of the University of Bern. The C storage per plot is the difference between the percentage carbon in autumn 2017 and in autumn 2015.

### Analyses

#### Multifunctionality

The functions we measured are all related to fast cycling and low resource losses, and thus we considered all to be positive when calculating our “ecosystem function multifunctionality” metrics (Manning et al., 2018). We used a multiple threshold approach because, unlike the averaging approach, it does not assume substitutability of functions and can distinguish cases where functions are at intermediate values from cases where some functions are high and other low (see Byrnes et al., 2014; Manning et al., 2018). We transformed each function in order to fit more closely to a normal distribution, which means functions contribute more equally to multifunctionality (see Table S9 and Figure S6) and standardised the data around the mean to have all functions on a common scale (using the formula x-mean/sd). We then calculated multifunctionality using thresholds from 0.5 to 0.8 with an interval of 0.05 (Le Bagousse-Pinguet et al., 2019). We excluded very large (>0.8) or very small (<0.5) thresholds as multifunctionality values tend to have small ranges at the extremes (i.e. most plots either have 0 multifunctionality or maximum multifunctionality). The thresholds were defined relative to the mean of the five highest values of each function, in order to reduce the influence of outliers (Zavaleta et al., 2010). Multifunctionality was the proportion of functions reaching this threshold per plot, calculated using the multidiv function (available at https://github.com/eric-allan/multidiversity).

We then tested the effects of our experimental treatments on multifunctionality: N enrichment, fungicide, species richness, SLA (as a continuous measure of functional composition), MPD, and their interaction. We used sown species richness as our main measure, following recommendations and common practice in analysing biodiversity experiment data (Schmid et al., 2002) (however, using realised richness resulted in very similar results, see Table S4). However, we used realised CWM SLA because functional composition should be important if mass ratio effects (Grime, 1998) dominate and weighting trait values by abundance better reflects the theoretical expectation. We explored both realised and abundance weighted effects of functional diversity (MPD in SLA), see above. We did not test for SLA x MPD interactions, as the two factors are not fully orthogonal in the experimental design (MPD is inevitably maximal at intermediate SLA, see Figure S5). We allowed all possible interactions between the other experimental treatments.

The effects of these different factors can vary according to the threshold used to calculate multifunctionality. Previous approaches have tested how diversity effects vary depending on the threshold (Byrnes et al., 2014) but visualising how multiple factors vary with multifunctionality thresholds is more challenging. We therefore used a new approach to test which treatment effects were consistent across thresholds and which depended on the threshold level. To do this, we first calculated multifunctionality per plot at seven different thresholds: 50, 55, 60, 65, 70, 75 and 80% thresholds. We then analysed all the multifunctionality values using linear mixed effects models, fitting all the treatment variables described above as fixed effects. We included *plot number*, *block* and *species combination* (random assemblages within diversity and functional composition levels) as random terms. We also included threshold as a fixed continuous term which could interact with the treatment variables, and random slopes for the effect of threshold on multifunctionality in each plot, which corrects for additional autocorrelation between multifunctionality values measured in the same plot. Our data met the model assumptions of homogeneity of variance and normal distribution of residuals. All explanatory variables were scaled and centred to a mean of 0 and standard deviation of 1, so that their effect sizes are comparable. Fitting all multifunctionality measures in the same model is a similar approach to that used in Soliveres, van der Plas, et al. (2016) but extended to multifunctionality calculated at different thresholds. Using this method, we could estimate which effects were significant across all thresholds, and which ones changed continuously as the threshold was increased. We then estimated the overall effect of each treatment and interaction across the thresholds and calculated a confidence interval around the effect. Confidence intervals were calculated using a script from Ben Bolker (available at https://bbolker.github.io/mixedmodels-misc/glmmFAQ.html#predictions-andor-confidence-or-prediction-intervals-on-predictions).

Since any multifunctionality metric depends on the number and identity of functions, which makes generalisation across studies challenging (Bradford et al., 2014; Meyer et al., 2018; Giling et al., 2019; Jing et al., 2020), all functions were also analysed separately using linear mixed effect models with the same fixed and random terms.

Monocultures all get an arbitrary MPD of 0 causing some correlation between MPD and species richness (Figure S5). To ensure that the effect of diversity was not driven only by the difference between monocultures and polycultures, we did an additional analysis in which we added a binary variable for the monoculture/not monoculture contrast. Results using this approach were qualitatively similar to those presented in the main text (Table S15). Replacing the effect of sown by realised species richness gave similar results (Table S4).

All analyses were conducted in R using the package lme4 (Bates et al., 2015; R. Core Team, 2019). We simplified full models by dropping terms that did not significantly improve the overall model fit using likelihood-ratios. All models had conditional R-squared higher than 0.87.

#### Effects of individual species on functions

To better understand the interaction between species richness and SLA, we conducted an additional analysis testing if individual species from the same functional group (slow or fast) were similar in the set of functions they increased or decreased, and if there were trade-offs between the functions supplied by different species of the same group. In short, we followed a six-step approach: i) extract the functional effect of each species using linear models (Hector & Bagchi, 2007; Isbell et al., 2011), ii) calculate the similarity in these functional effects in all possible 4-species assemblages, iii) repeat step ii for both positive and negative functional effects, iv) analyse if that similarity in functional effects changes depending on the mean (and variation in) SLA of the 4-species assemblages. If low-SLA assemblages show higher similarity, this means that slow-growing species are more functionally redundant than fast-growing species. Finally, two more steps were performed to analyse the likelihood that species with similar functional (slow-growing) trait values showed stronger functional trade-offs: steps v) take all pairwise species combinations and quantify the number of times that one of the species in the pair have a functional effect of opposite sign than the other species in the pair (e.g., one has a positive effect on biomass production and the other a negative effect on the same function), and vi) relate the number of functional trade-offs (as quantified in step v) to the mean and difference in SLA between the two species in the pair. If the number of trade-offs was higher in pairs of species with low SLA, then we can conclude than slow-growing species are more likely to have opposite functional effects than may cancel each other out.

First step: we tested the effect of each species on each function, following approaches in Hector & Bagchi (2007) and Isbell et al. (2011). To do so, we first extracted the effect of species on each function. We fitted two separate models, one with the presence and one with the abundance (relative cover) of each of the 20 species, using the following model structure:

*Function ∼ Nitrogen * Fungicide + Species1 + … + Species20 + (1|Block)*

Non-significant effects were set to 0 (Table S7 and Table S8).

We then calculated (step ii) how similar different sets of species were in the functions they affected in order to test how many functions different communities of fast vs. slow species would be expected to provide. We calculated the similarity in functional effects for all possible combinations of four species. We chose four species as this allows us to test the effect of different species combinations, varying in mean and variance of their functional traits, and because it corresponds to one of our diversity treatments. However, we also tested combinations of 3 and 6 species, which did not change the results (Table S6). We calculated the similarity in both positive and negative functional effects for all combinations of four species (step iii), and for effects of species presence and abundance. We used the CqN similarity index (Jost et al., 2011), which is a common measure of beta diversity and allows a calculation of dissimilarity among any number of communities. We weighted the effects to give more importance to larger functional effects by using a q value of 2. This corresponds to the multicommunity extension of the Morisita-Horn index.

We then calculated the mean and variance in SLA between all sets of four species and regressed them against the dissimilarity in functional effects between the four species (CqN; step iv). We were principally interested in the effect of mean SLA, to determine if sets of species with higher mean SLA are more different in the functions they provide. However, we included the variance in SLA between the species to account for the fact that a set of four species with intermediate mean SLA could either be composed of low and high SLA (slow and fast) species or a set of species all with intermediate SLA. To test if the effects were significant, we randomly shuffled the SLA values between the species and recalculated the relationships between mean and variance in SLA and functional dissimilarity 1000 times. We then tested whether the observed slopes between mean SLA and variance in SLA and functional dissimilarity (CqN) lay outside the 2.5 and 97.5% quantiles of the random slopes. These analyses were repeated for the positive and negative effects and effects of species presence and abundance, see Table S6.

Then, we tested for trade-offs between the functions supplied by different species and whether this was related to their SLA (steps v and vi). For each pair of species, we summed the number of significant functional effects that were opposing (e.g. one species has a positive effect on a given function while the other species has a negative effect). We related the number of trade-offs to the mean and difference in SLA for the species pair, to test if pairs of low SLA (slow) species tend to have more opposing effects. We again randomly shuffled the SLA values between the species, recalculated the relationships 1000 times and tested whether the observed slopes lay outside the 2.5 and 97.5% quantiles of the random slopes, see Table S6.

#### Number and identity of functions

In addition to the threshold chosen, multifunctionality values depend on the number and identity of functions included, which makes generalisation across studies challenging (Bradford et al., 2014; Meyer et al., 2018; Giling et al., 2019; Jing et al., 2020). To test the sensitivity of our results to the number and identity of functions, we calculated multifunctionality for all possible combinations of functions between two and ten functions (1013 combinations), and for four thresholds from 0.5 to 0.8 (4052 multifunctionality measures). We extracted the effects of our treatments and their interactions from a linear mixed effect model fitted to each multifunctionality measure. In order to correct for the uncertainty in each estimate, we weighted each estimate by the inverse of its standard error. We then calculated an overall mean and confidence interval for multifunctionality per threshold depending on the number of functions included. The number of functions included did not significantly change the mean effect of our variables on multifunctionality, except for the interaction between SLA and species richness at the 50% threshold, which strengthened as more functions were considered. The variation around the mean got larger with a decreasing number of functions, going from positive to negative depending on the functions included (Figure S2 and Figure S3). This variation was driven more by variation in the identity rather than in the number of functions included (Figure S2).

## Acknowledgements

We would like to thank Pete Manning and Jasper van Ruijven for very helpful comments on earlier drafts of the manuscript. We would also like to thank Mervi Laitinen and Marlise Zimmermann, the technicians working with us on the PaNDiv Experiment, as well as the large team of helpers on the field and in the lab. This study was supported by funding of the Swiss National Science Foundation (31003A_160212). SS was supported by the Spanish Government under a Ramón y Cajal contract (RYC- 2016- 20604).

## Authors contribution

N.A.P., S.L.C. and E.A. designed and set up the PaNDiv Experiment; N.A.P., S.L.C., T.M., TZ.N., H.V. collected the data; N.A.P., T.M., N.H., V.H.K. and T.K. processed and analysed the NIRS samples; N.A.P. analysed the data and wrote the first manuscript with substantial input from E.A., S.S. and S.L.C. All authors contributed to revisions of the manuscript.

