## Supplementary information for "Nitrogen availability and plant functional composition modify biodiversity-multifunctionality relationships"

**Table S1: PCA axis models.** Effect on multifunctionality of sown species richness (SR), functional diversity (MPD), functional composition (PCA1), fungicide (Fng), nitrogen enrichment (N), multifunctionality threshold and their interaction. Output of the linear mixed effects model after model simplification (scaled variables).

In these models, we replaced community weighted mean specific leaf area (CWM SLA) by the first axis of a principal component analysis (PCA).

Initial model:
multifunctionality ~ Threshold * (N + PCA1 + Fng + SR + MPD) ^2
- Threshold: PCA1:MPD - PCA1:MPD + (1|Block) + (1|Combination) + (Threshold |Plot_Nr)

a) PCA including CWM SLA and CWM LDMC (leaf dry matter content). Functional diversity (MPD) is calculated using both SLA and LDMC.

Importance of components:

| | PC1 | PC2 |
|---|---|---|
| Standard deviation | 1.07 | 0.93 |
| Proportion of Variance | 0.57 | 0.43 |
| Cumulative Proportion | 0.57 | 1.00 |

Correlation CWM SLA and PC1: **-0.76**

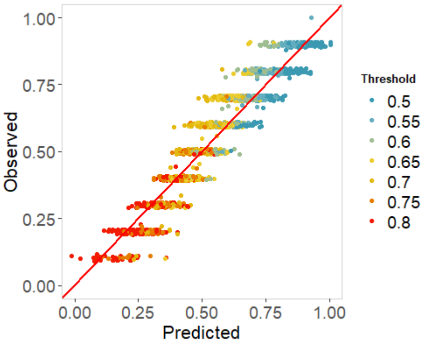


| Factor | Estimate | Std.Error | Significance |
|---|---|---|---|
| (Intercept) | 0.568 | 0.015 | |
| Threshold | -0.179 | 0.004 | <0.001 |
| N | 0.014 | 0.005 | |
| Fng | -0.009 | 0.005 | |
| SR | 0.000 | 0.012 | |
| PC1 | -0.033 | 0.010 | |
| MPD | 0.005 | 0.009 | |
| **N x MPD** | -0.010 | 0.004 | 0.018 |
| **Fng x MPD** | -0.008 | 0.004 | 0.045 |
| **SR x PC1** | -0.021 | 0.010 | 0.033 |

Conditional R-squared: 0.88

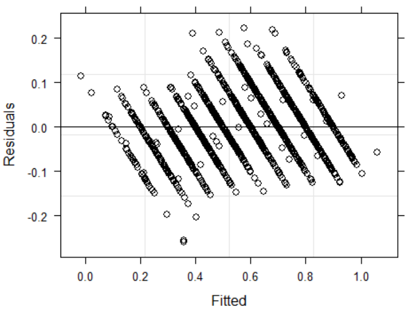


Relationship between SLA and PC1 per species.

Species list, see Table S11.

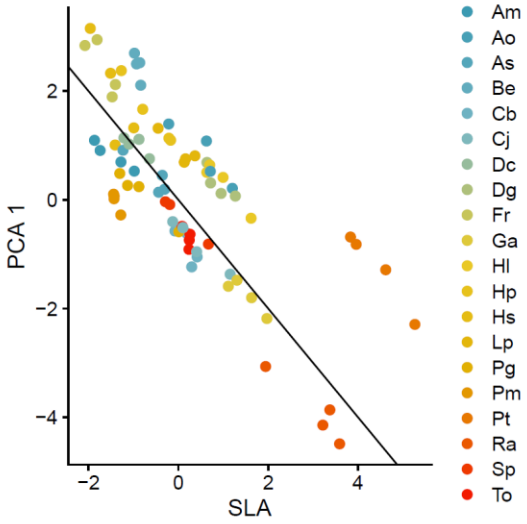

b) PCA including CWM SLA, CWM LDMC and CWM biomass nitrogen. Functional diversity (MPD) is calculated using SLA, LDMC, and biomass nitrogen concentration.

Importance of components:

| | PC1 | PC2 | PC3 |
|---|---|---|---|
| Standard deviation | 1.08 | 1.04 | 0.86 |
| Proportion of Variance | 0.39 | 0.36 | 0.25 |
| Cumulative Proportion | 0.39 | 0.75 | 1.00 |

Correlation CWM SLA and PC1: **0.84**

| Factor | Estimate | Std.Error | Significance |
|---|---|---|---|
| (Intercept) | 0.575 | 0.015 | |
| Threshold | -0.179 | 0.004 | <0.001 |
| N | 0.013 | 0.005 | |
| SR | 0.010 | 0.012 | |
| PC1 | 0.029 | 0.008 | |
| MPD | 0.002 | 0.009 | |
| **N x MPD** | -0.010 | 0.004 | 0.010 |
| **SR x PC1** | 0.031 | 0.007 | <0.001 |

Conditional R-squared: 0.88

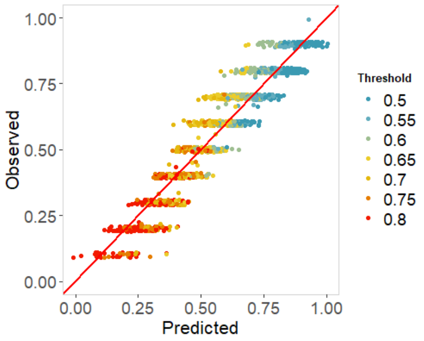


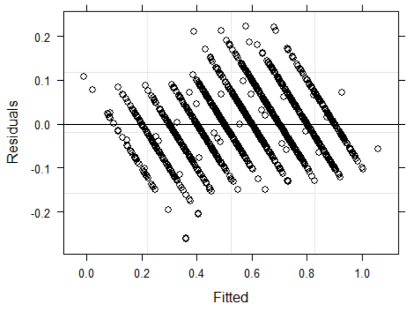


Relationship between SLA and PC1 per species.

Species list, see Table S11.

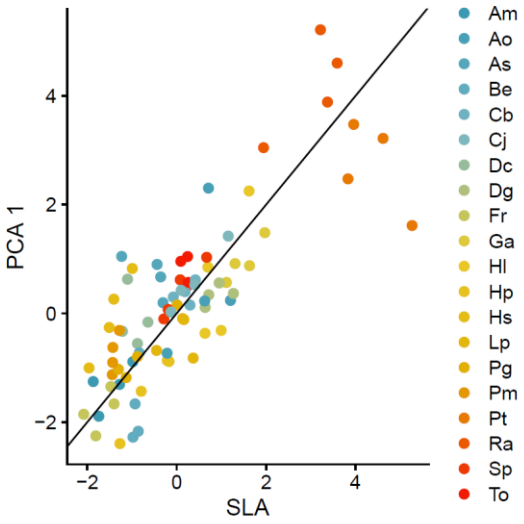

**Table S2: Main model.** Effect on multifunctionality of species richness (SR), functional diversity (MPD), community weighted mean specific leaf area (SLA), fungicide (Fng), nitrogen enrichment (N), multifunctionality threshold and their interaction. Output of the linear mixed effects model after model simplification (scaled variables).

Initial model in R:

multifunctionality ~
Threshold * (N + SLA + Fng + SR + MPD) ^2
- Threshold:SLA:MPD - SLA:MPD
+ (1|Block) + (1|Combination) + (Threshold|Plot_Nr)

| Factor | Estimate | Std.Error | Significance |
|---|---|---|---|
| (Intercept) | 0.572 | 0.014 | |
| Threshold | -0.179 | 0.004 | <0.001 |
| N | 0.014 | 0.005 | |
| Fng | -0.008 | 0.004 | |
| SR | 0.010 | 0.011 | |
| SLA | 0.033 | 0.009 | |
| MPD | 0.001 | 0.009 | |
| **N x MPD** | -0.010 | 0.004 | 0.014 |
| **Fng x MPD** | -0.008 | 0.004 | 0.039 |
| **SR x SLA** | 0.025 | 0.009 | 0.005 |

Conditional R-squared: 0.88

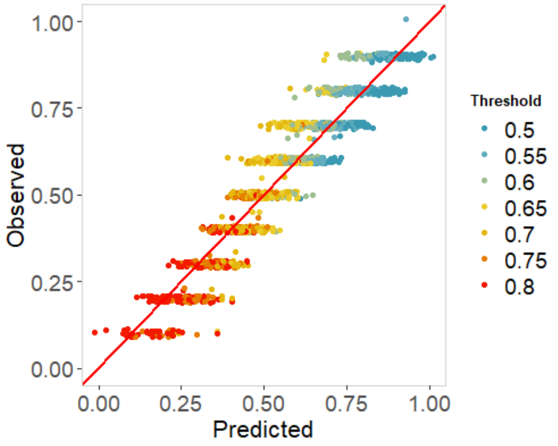


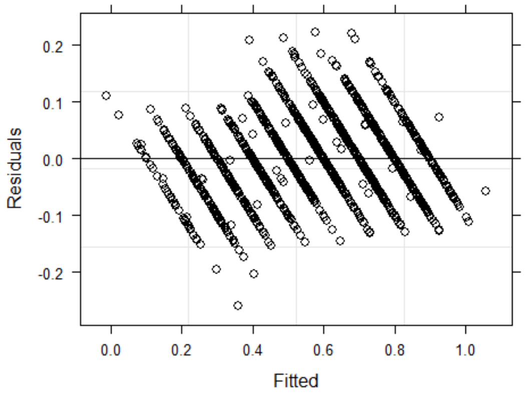

**Table S3: No phosphatase model.** Effect on multifunctionality, calculated without phosphatase activity, of sown species richness (SR), functional diversity (MPD), community weighted mean specific leaf area (SLA), fungicide (Fng), nitrogen enrichment (N), multifunctionality threshold and their interaction. Output of the linear mixed effects model after model simplification (scaled variables).

Initial model:
multifunctionality ~
Threshold * (N + SLA + Fng + SR + MPD) ^2
- Threshold:SLA:MPD - SLA:MPD + (1|Block) + (1|Combination) + (Threshold |Plot_Nr)

| Factor | Estimate | Std.Error | Significance |
|---|---|---|---|
| (Intercept) | 0.554 | 0.015 | |
| Threshold | -0.172 | 0.004 | <0.001 |
| N | 0.014 | 0.005 | |
| SR | 0.008 | 0.013 | |
| SLA | 0.035 | 0.010 | |
| MPD | 0.001 | 0.009 | |
| **N x MPD** | -0.008 | 0.004 | 0.047 |
| **SR x SLA** | 0.030 | 0.010 | 0.002 |

Conditional R-squared: 0.87

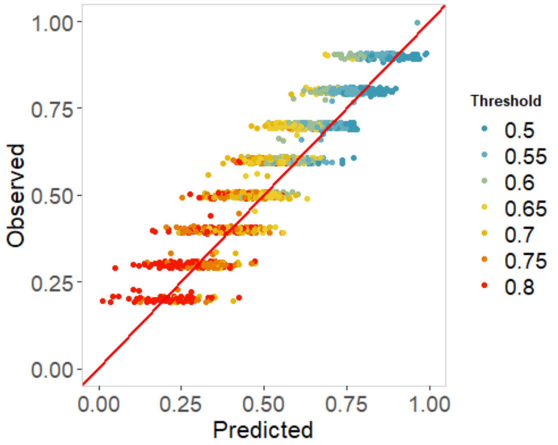


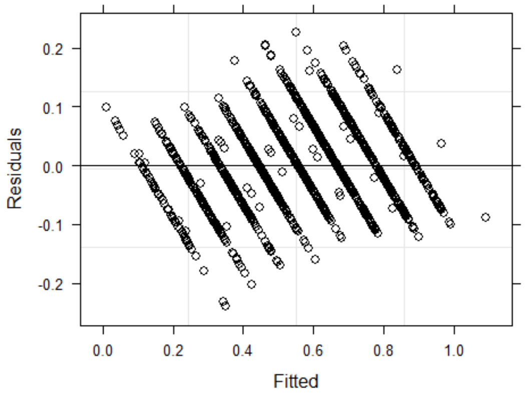

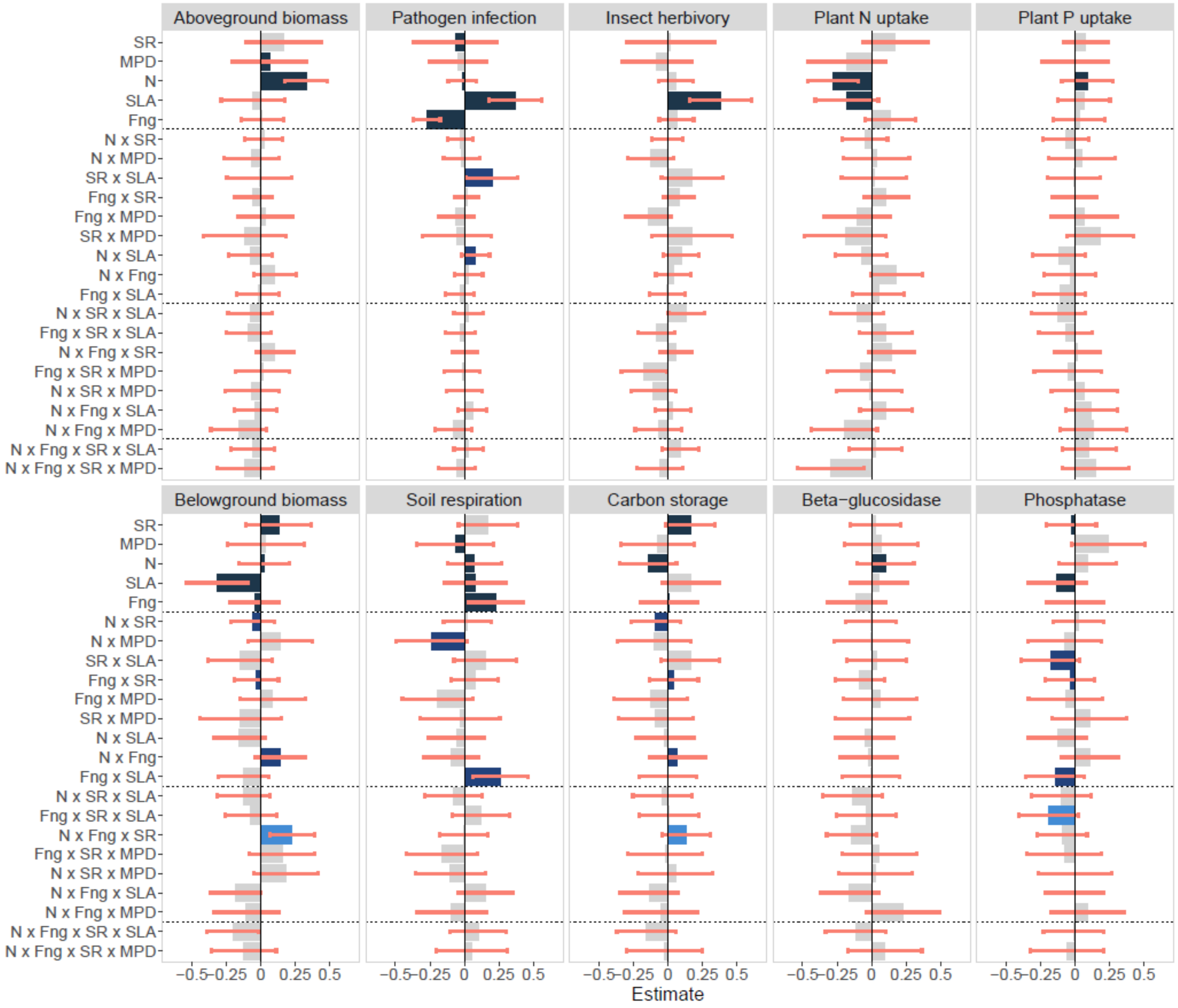


**Figure S1:** Drivers of individual functions. Mean standardised effect (+/- confidence intervals) of all variables and their interaction on functions. Non-significant effects are shown in grey. Effects and confidence intervals are extracted from the full model, while significance was assessed after model simplification and after dropping the higher order terms (i.e. significance of two way interactions was assessed after three way interactions were removed), so error bars for marginal terms do not necessarily correspond to their significance.

SR: sown plant species richness, MPD: mean pairwise distance in SLA equally weighted, N: nitrogen enrichment, SLA: abundance weighted community specific leaf area, Fng: fungicide treatment. Each function was measured on 216 plots.

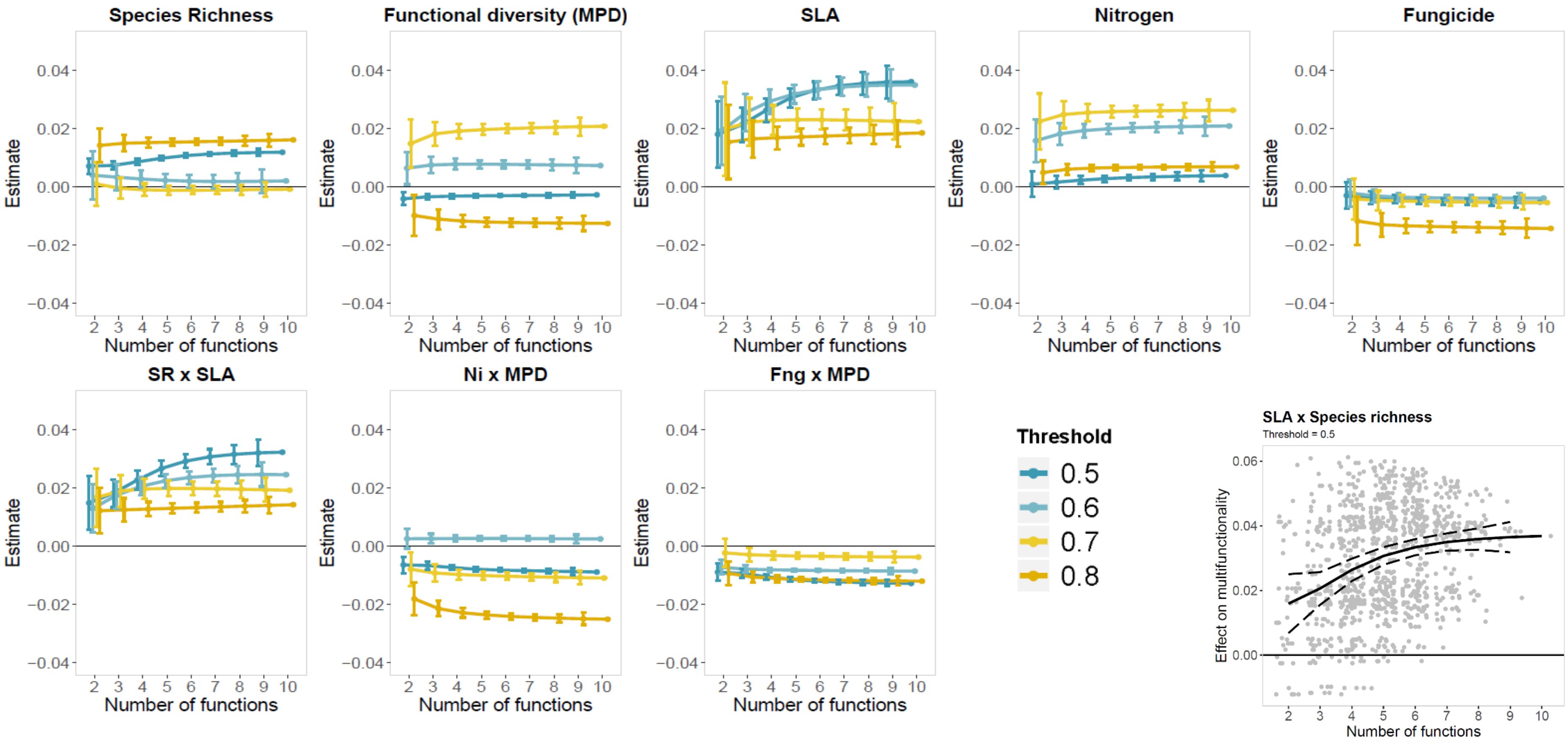


**Figure S2:** Estimate (+/- confidence interval) of manipulated variables and interactions effect on multifunctionality, depending on the threshold and the number of functions. Mean effect (+/- CI) of the species richness x SLA interaction on multifunctionality at a 0.5 threshold depending on the number of functions. In grey are shown the transformed estimates for each combination of function.

Effects were extracted from linear mixed effects models and weighted by the inverse of the square root of their standard error. We then calculated mean and CI per number of functions and per threshold, see methods

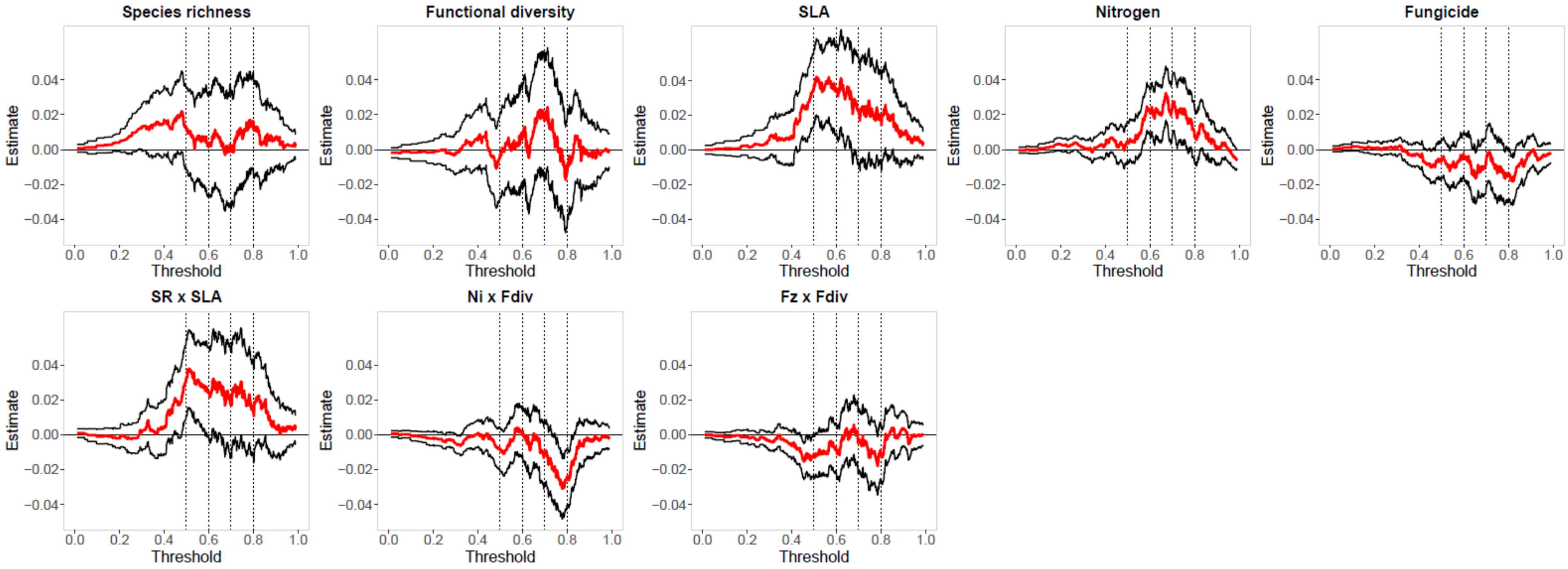


**Figure S3:** Estimate (+/- confidence interval) of manipulated variables and interactions effect on multifunctionality, depending on the thresholds. Multifunctionality was calculated using all ten functions.

Effects were extracted from linear mixed effects models and weighted by the inverse of the square root of their standard error. We then calculated mean and CI per threshold, see methods.

**Table S4: Realised diversities models.** Effect on multifunctionality of realised species richness (RSR), functional diversity (MPD), community weighted mean specific leaf area (SLA), fungicide (Fng), nitrogen enrichment (N), multifunctionality threshold and their interaction. Output of the linear mixed effects model after model simplification (scaled variables). Realised species richness includes per plot all species that were recorded at least once between 2017 and 2018.

Initial model:
multifunctionality ~ Threshold * (N + SLA + Fng + RSR + MPD) ^2
- Threshold:SLA:MPD - SLA:MPD + (1|Block) + (1|Combination) + (Threshold |Plot_Nr)

| Factor | Estimate | Std.Error | Significance |
|---|---|---|---|
| (Intercept) | 0.571 | 0.014 | |
| Threshold | -0.179 | 0.004 | <0.001 |
| N | 0.014 | 0.005 | |
| Fng | -0.008 | 0.004 | |
| RSR | 0.004 | 0.013 | |
| SLA | 0.030 | 0.009 | |
| MPD | 0.004 | 0.009 | |
| **N x MPD** | -0.010 | 0.004 | 0.018 |
| **Fng x MPD** | -0.008 | 0.004 | 0.043 |
| **RSR x SLA** | 0.028 | 0.011 | 0.008 |

Conditional R-squared: 0.88

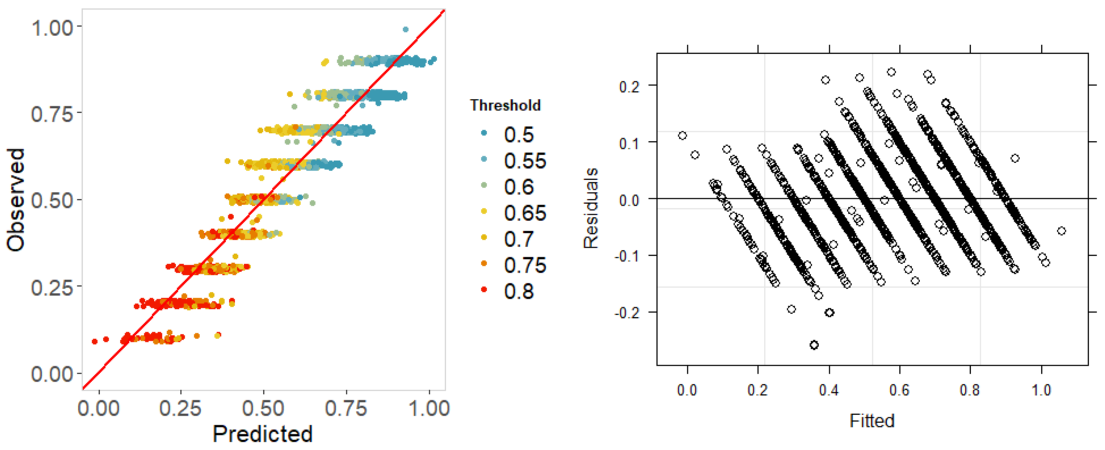

Effect on multifunctionality of species richness (SR), equally weighted realised functional diversity (RMPD), community weighted mean specific leaf area (SLA), fungicide (Fng), nitrogen enrichment (N), multifunctionality threshold and their interaction. Output of the linear mixed effects model after model simplification (scaled variables). Realised functional diversity includes per plot all species that were recorded at least once between 2017 and 2018. RMPD x Fng is marginally significant (pvalue 0.057).

Initial model:
multifunctionality ~ Threshold * (N + SLA + Fng + SR + RMPD) ^2
- Threshold:SLA:RMPD - SLA:RMPD + (1|Block) + (1|Combination) + (Threshold |Plot_Nr)

| Factor | Estimate | Std.Error | Significance |
|---|---|---|---|
| (Intercept) | 0.571 | 0.015 | |
| Threshold | -0.180 | 0.004 | <0.001 |
| N | 0.013 | 0.005 | |
| SR | 0.012 | 0.011 | |
| SLA | 0.033 | 0.009 | |
| RMPD | -0.002 | 0.008 | |
| Fng | -0.008 | 0.004 | |
| **N x RMPD** | -0.011 | 0.004 | 0.038 |
| **RMPD x Fng** | -0.008 | 0.004 | 0.057 |
| **SR x SLA** | 0.025 | 0.009 | 0.007 |

Conditional R-squared: 0.88

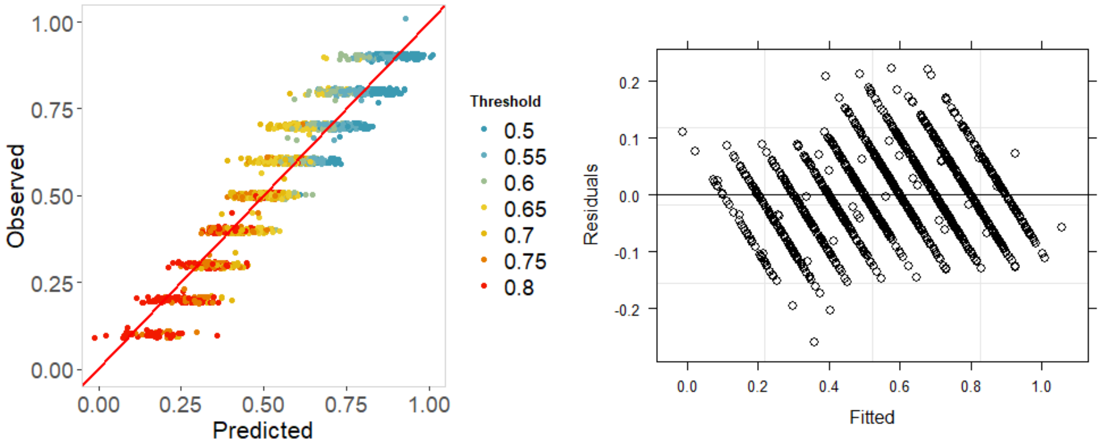

**Table S5: No pathogens model.** Effect on multifunctionality, calculated without pathogen infection, of sown species richness (SR), functional diversity (MPD), community weighted mean specific leaf area (SLA), fungicide (Fng), nitrogen enrichment (N), multifunctionality threshold and their interaction. Output of the linear mixed effects model after model simplification (scaled variables).

Initial model:
multifunctionality ~
Threshold * (N + SLA + Fng + SR + MPD) ^2
- Threshold:SLA:MPD - SLA:MPD + (1|Block) + (1|Combination) + (Threshold |Plot_Nr)

| Factor | Estimate | Std.Error | Significance |
|---|---|---|---|
| (Intercept) | 0.586 | 0.016 | |
| Threshold | -0.182 | 0.004 | <0.001 |
| N | 0.020 | 0.005 | |
| SR | 0.012 | 0.012 | |
| MPD | 0.005 | 0.009 | |
| **Threshold x SR** | 0.009 | 0.004 | 0.022 |
| **Threshold x MPD** | -0.009 | 0.004 | 0.027 |

Conditional R-squared: 0.87

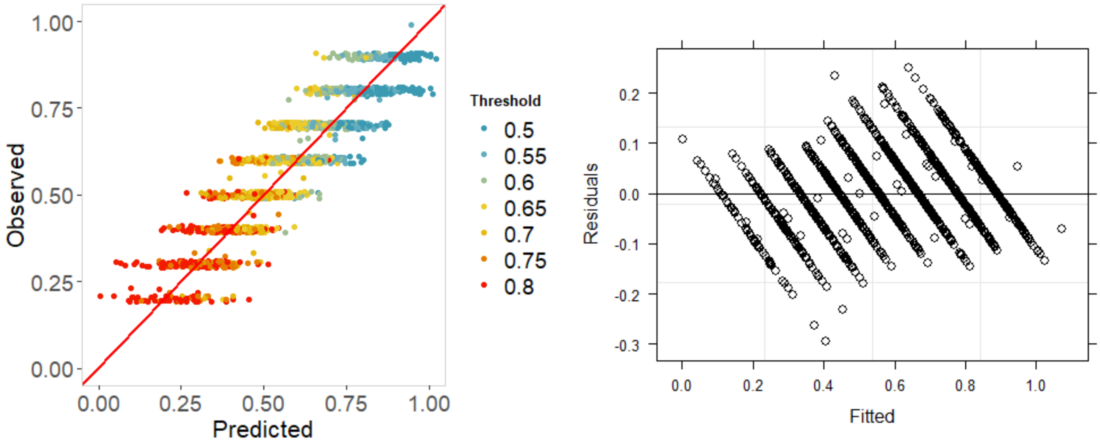

**Table S6:** Correlations between the mean and variance in SLA for a set of four species and the similarity in the functional effects of the constituent species. The results were similar for sets of three and six species. For calculations details, see methods. We randomly shuffled the SLA values between the species and recalculated the relationships 1000 times. The 2.5% and 97.5% quantiles for the distribution of random slopes are shown, when the observed slope lies outside this range we consider there to be a significant effect of mean/variance in SLA on functional dissimilarity.

| **SIMILARITY** | | | **Model slope** | **<0.025** | **>0.975** |
|---|---|---|---|---|---|
| **Presence** | | | | | |
| | **Positive** | Mean SLA | -7.15E-03 | -3.35E-03 | 3.33E-03 |
| | | Variance SLA | -5.84E-04 | -3.15E-04 | 3.44E-04 |
| | **Negative** | Mean SLA | -9.04E-03 | -4.50E-03 | 4.69E-03 |
| | | Variance SLA | 8.74E-04 | -3.43E-04 | 3.50E-04 |
| **Abundance** | | | | | |
| | **Positive** | Mean SLA | -3.94E-02 | -1.19E-02 | 1.22E-02 |
| | | Variance SLA | -1.49E-03 | -1.01E-03 | 9.40E-04 |
| | **Negative** | Mean SLA | -7.36E-03 | -2.10E-03 | 2.07E-03 |
| | | Variance SLA | 7.45E-04 | -1.61E-04 | 1.58E-04 |

Effect of SLA on the number of trade-offs between positive and negative effects on function for each pair of species.

| **TRADE-OFFS** | | **Model slope** | **<0.025** | **>0.975** |
|---|---|---|---|---|
| **Presence** | Mean SLA | 2.93E-03 ns | -1.94E-02 | 2.06E-02 |
| | Variance SLA | 1.53E-03 | -1.34E-03 | 1.39E-03 |
| **Abundance** | Mean SLA | -1.80E-02 ns | -2.21E-02 | 2.26E-02 |
| | Variance SLA | 1.85E-03 | -1.31E-03 | 1.43E-03 |

**Table S7:** Mean effect of individual species presence on individual functions. Positive effects are coloured in blue, negative in red. For calculations details, see methods.

| | | Aboveground functions | | | | | Belowground functions | | | | |
|---|---|---|---|---|---|---|---|---|---|---|---|
| | | Aboveground biomass | Pathogen infection | Insect herbivory | Plant N uptake | Plant P uptake | Belowground biomass | Soil respiration | Carbon storage | β -glucosidase | Phosphatase |
| fast | As | | | | | | | | | | |
| | Cb | | | | 0.524 | | | | | | |
| | Dg | | 0.277 | | | | | | | | |
| | Ga | 0.721 | | -0.715 | -0.471 | | | | | | |
| | Hl | | 0.451 | | | | | | | | |
| | Hs | -0.643 | 0.454 | | | | -0.498 | | | | |
| | Lp | | 0.717 | | | | | | | | |
| | Pt | | -0.627 | | -0.655 | | | | | | |
| | Ra | | | 0.809 | | | -0.585 | | | | |
| | To | | | | | | | | | | |
| slow | Am | | -0.595 | -1.426 | 0.820 | | 0.548 | | | | |
| | Ao | -0.670 | | | | | -0.890 | | | | |
| | Be | | 0.638 | | | | | | | | |
| | Cj | 0.960 | 0.542 | 0.540 | | | | | | | |
| | Dc | | -0.784 | | | | | | | | |
| | Fr | 0.508 | | | | | 0.499 | 0.510 | | | |
| | Hp | | | | | | | -0.496 | | | |
| | Pg | | | -0.422 | | | 0.494 | | | | |
| | Pm | | -0.860 | | | | | | | | |
| | Sp | | -0.466 | 0.883 | 0.627 | | | | | | |

**Table S8:** Mean effect of individual species, abundance weighted, on individual functions. Positive effects are coloured in blue, negative in red. For calculations details, see methods.

| | | Aboveground functions | | | | | Belowground functions | | | | |
|---|---|---|---|---|---|---|---|---|---|---|---|
| | | Aboveground biomass | Pathogen infection | Insect herbivory | Plant N uptake | Plant P uptake | Belowground biomass | Soil respiration | Carbon storage | β -glucosidase | Phosphatase |
| fast | As | | -1.025 | -1.718 | | | | | | -0.260 | |
| | Cb | | | -0.817 | | | | | | | |
| | Dg | | | | | | | -1.203 | | -0.278 | |
| | Ga | 1.449 | | -1.941 | | | | | | | |
| | Hl | | | -1.109 | -1.180 | | | | | | |
| | Hs | -1.520 | -0.679 | | | | | | | | |
| | Lp | | | -1.192 | | | | | | | |
| | Pt | -0.868 | -2.344 | -0.693 | -1.451 | | | | | | |
| | Ra | -1.167 | -0.964 | 1.872 | | | | | | | |
| | To | | | | | | | | | | |
| slow | Am | 1.128 | -2.165 | -3.001 | 1.370 | | | | | | |
| | Ao | | -1.332 | -1.505 | | | | | | | |
| | Be | | | | | | | | | -0.355 | |
| | Cj | 1.976 | -0.770 | | | | | | | | |
| | Dc | -1.405 | -2.848 | -2.142 | | | | -1.650 | | -0.253 | |
| | Fr | 1.096 | -1.326 | -1.685 | | | | | | | |
| | Hp | | | -1.131 | | | | -1.154 | | | |
| | Pg | | -0.921 | -1.620 | | | | | | -0.233 | |
| | Pm | | -3.354 | | | | | | | | |
| | Sp | | | | | | | | | | |

**Figure S4: Raw data plot of the main text Figure 2b.** Effect of specific leaf area (SLA, $m^2\ kg^{-1}$) and species richness on multifunctionality, at a 65% threshold.

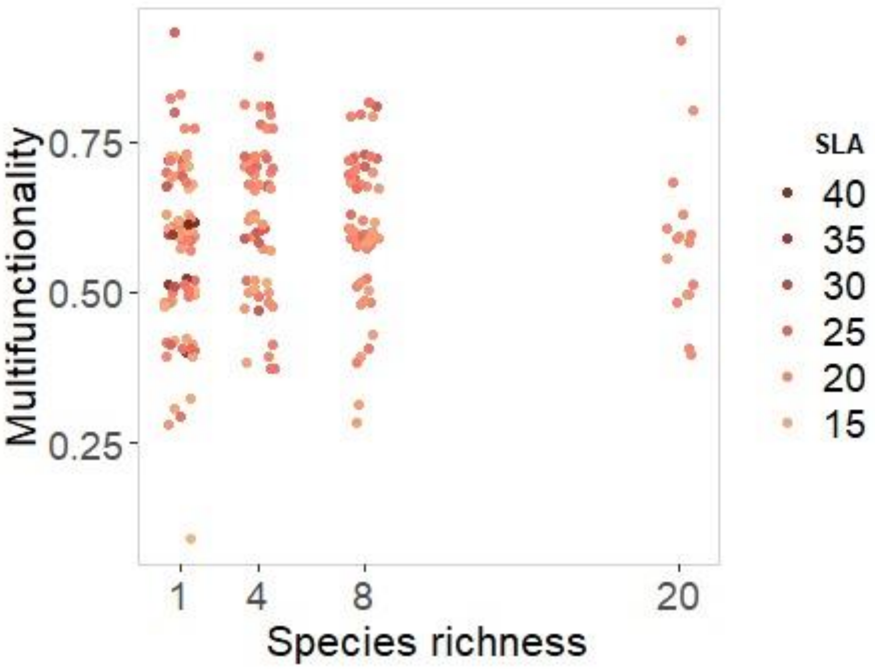

**Table S9:** Description of the functions included in our multifunctionality measure.

| Variable / Measurement | Number of measurements | Time of measurement | Related function | Transformation |
|---|---|---|---|---|
| **Aboveground biomass** | 4 | June & Aug 17, June & Aug 18 | Aboveground primary production | Sqrt |
| **Pathogen infection** | 3 | Oct 17, July & Oct 18 | Transfer of energy to higher trophic levels | - |
| **Insect herbivory** | 2 | June & Aug 18 | Transfer of energy to higher trophic levels | Sqrt |
| **Plant N uptake** | 2 (plant), 4 (soil) | June & Aug 18 (plant), Apr, Mai, July & Aug 18 (soil) | N cycling, community N uptake efficiency | Log |
| **Plant P uptake** | 2 (plant), 4 (soil) | June & Aug 18 (plant), Apr, Mai, July & Aug 18 (soil) | P cycling, community P uptake efficiency | Log |
| **Belowground biomass** | 1 | Autumn 17 | Belowground primary production | Log |
| **Soil respiration** | 5 | Mai to Sept 2018 | C cycling, microbial activity | Log |
| **Carbon storage** | 1 | Autumn 2015 and 2017 | C cycling | Sqrt +1 |
| **β -glucosidase** | 5 | Autumn 17, Apr, Mai, July & Aug 18 | C cycling, microbial activity | Sqrt |
| **Phosphatase** | 5 | Autumn 17, Apr, Mai, July & Aug 18 | P cycling, microbial activity | Log |
| **Plant % cover** | 4 | June & Aug 17, June & Aug 18 | | |
| **Plant traits** | 3 | Aug 17, June & Aug 18 | | |

**Table S10: Weighted MPD model.** Effect on multifunctionality of species richness (SR), abundance weighted functional diversity (MPDw), community weighted mean specific leaf area (SLA), fungicide (Fng), nitrogen enrichment (N), multifunctionality threshold and their interaction. Output of the linear mixed effects model after model simplification (scaled variables).

Initial model in R:

multifunctionality ~
Threshold * (N + SLA + Fng + SR + MPDw) ^2
- Threshold:SLA:MPDw - SLA:MPDw
+ (1|Block) + (1|Combination) + (Threshold|Plot_Nr)

| Factor | Estimate | Std. Error | Significance |
|---|---|---|---|
| (Intercept) | 0.589 | 0.016 | |
| Threshold | -0.179 | 0.004 | <0.001 |
| N | 0.014 | 0.005 | |
| SR | 0.015 | 0.012 | |
| SLA | 0.036 | 0.009 | |
| MPDw | -0.013 | 0.010 | |
| **N x MPDw** | -0.011 | 0.004 | 0.009 |
| **SR x SLA** | 0.028 | 0.009 | 0.002 |
| **SR x MPDw** | -0.027 | 0.011 | 0.012 |

Conditional R-squared: 0.88

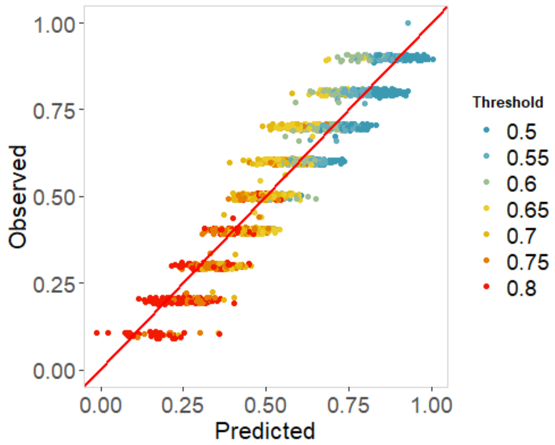


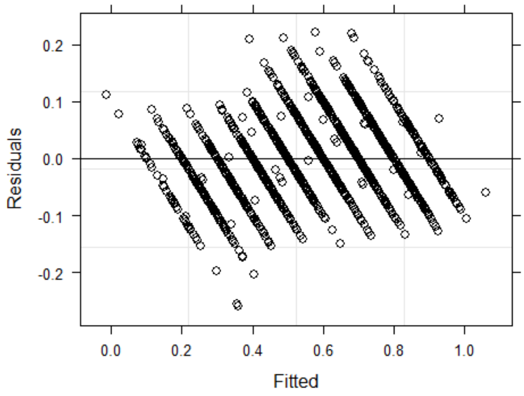

**Table S11:** PaNDiv Experiment list of species and corresponding growth strategy.

| Grasses | Leaf economics spectrum | Herbs | Leaf economics spectrum |
|---|---|---|---|
| *Poa trivialis*<br>*Lolium perenne*<br>*Holcus lanatus*<br>*Dactylis glomerata* | Fast | *Crepis biennis*<br>*Taraxacum officinale*<br>*Anthriscus sylvestris*<br>*Heracleum sphondylium*<br>*Galium album*<br>*Rumex acetosa* | Fast |
| *Helictotrichon pubescens*<br>*Festuca rubra*<br>*Bromus erectus*<br>*Anthoxanthum odoratum* | Slow | *Achillea millefolium*<br>*Centaurea jacea*<br>*Daucus carotta*<br>*Salvia pratensis*<br>*Prunella grandiflora*<br>*Plantago media* | Slow |

**Table S12: Soil pH does not respond to our different treatments.** Soil pH was measured in autumn 2017. Output of model simplification, after dropping terms that did not significantly improve the overall model fit using likelihood-ratios.

Initial model:
pH ~ N * Fng * SR * (SLA + MPD) + (1|Block) + (1|Combination)

| Factor | AIC | LRT | Pr(Chi) |
|---|---|---|---|
| <none> | -32.38 | <NA> | <NA> |
| N | -32.99 | 1.391 | 0.238 |
| Fng | -31.26 | 3.118 | 0.077 |
| N | -32.99 | 1.391 | 0.238 |
| SLA | -32.38 | 0.379 | 0.538 |
| SR | -30.76 | 0.339 | 0.560 |
| MPD | -29.1 | 0 | 0.999 |
| Fng x SR | -27.1 | 2.026 | 0.155 |
| N x Fng | -27.12 | 0.968 | 0.325 |
| N x SR | -26.09 | 0.013 | 0.911 |
| N x Fng x SR | -24.1 | 1.763 | 0.184 |
| Fng x SLA | -23.87 | 0.443 | 0.506 |
| N x SLA | -22.31 | 0.066 | 0.798 |
| N x Fng x SLA | -20.38 | 0.614 | 0.433 |
| Fng x MPD | -18.99 | 0.129 | 0.719 |
| SR x MPD | -17.12 | 0 | 0.984 |
| Fng x SR x MPD | -15.12 | 0.539 | 0.463 |
| SR x SLA | -13.66 | 0.011 | 0.916 |
| N x SR x SLA | -11.67 | 0.543 | 0.461 |
| Fng x SR x SLA | -10.21 | 0.437 | 0.509 |
| N x Fng x SR x SLA | -8.65 | 0.283 | 0.595 |
| N x MPD | -6.93 | 0.208 | 0.649 |
| N x SR x MPD | -5.14 | 0.245 | 0.621 |
| N x Fng x MPD | -3.38 | 0.001 | 0.971 |
| N x Fng x SR x MPD | -1.39 | 0.037 | 0.848 |

a)

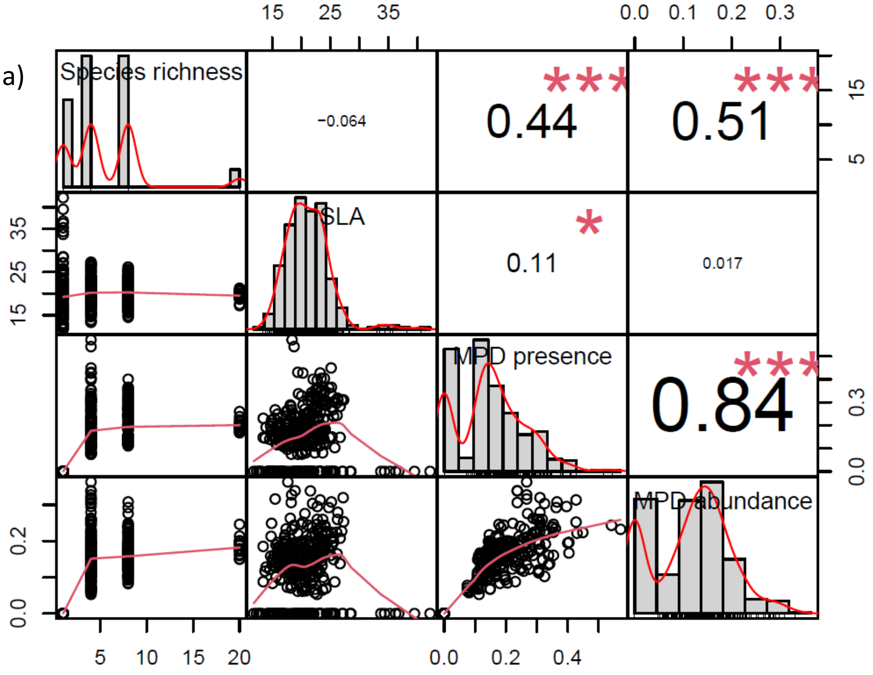


b)

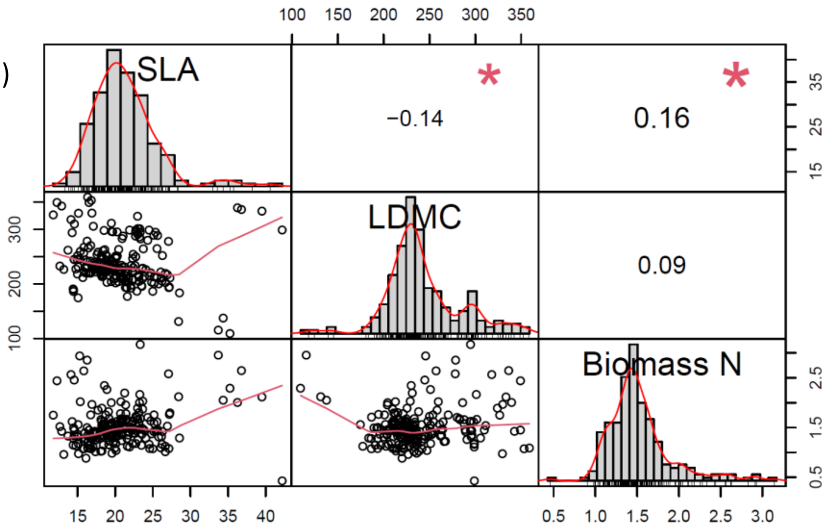


**Figure S5:**

a) Correlations between species richness, community weighted mean specific leaf area (SLA), functional diversity as mean pairwise distance based on species presence/absence (MPD presence) and based on species abundance (MPD abundance)

b) Correlations between community weighted mean specific leaf area (SLA), community weighted mean leaf dry matter content (LDMC) and biomass nitrogen content (Biomass N).

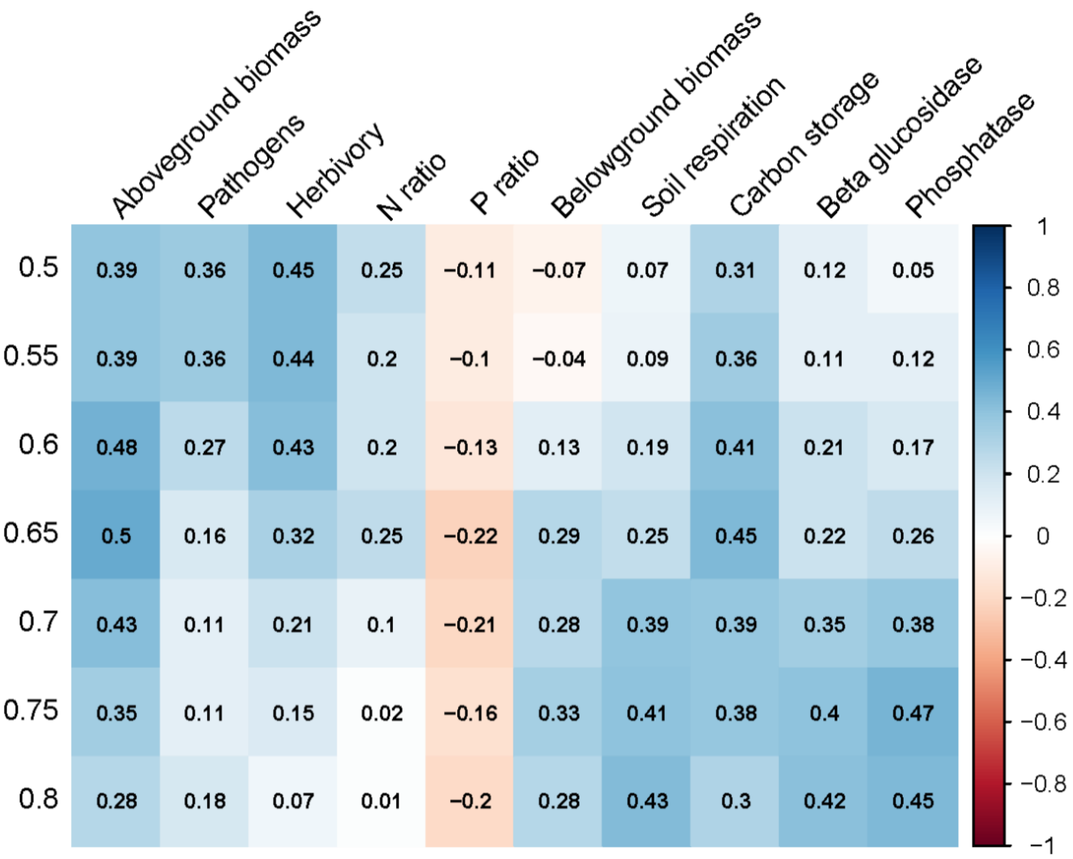


**Figure S6:** Pearson's correlations between the functions after normalisation (see Table S9 for the transformation detail) and multifunctionality at each threshold. Each function was measured on 216 plots. Although functions vary in how strongly correlated they are with multifunctionality there is no function that is very strongly correlated with the multifunctionality index. Responses of multifunctionality are therefore not driven by particular functions.

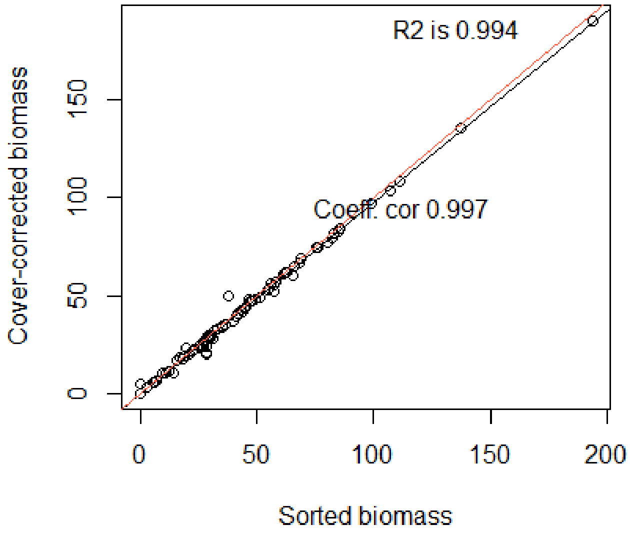


**Figure S7:** Correlation between biomass of target species based on sorted biomass and based on cover-corrected biomass.

From main text: The plot target biomass production (i.e. of the sown species without weeds) was calculated by multiplying the percentage cover of weeds by the total biomass and subtracting this estimated weed biomass from the total weight. To determine if the estimation was accurate, we sorted the biomass per species in June 2017 for all plots of one block. The sorted target biomass (without the weeds) was highly correlated with the estimated target biomass ($R^2$ = 0.994). The mean weed percentage was around five percent and therefore removing the weeds made little difference to the total biomass.

**Table S13: Bulk density does not respond to our different treatments.** Bulk density was measured in autumn 2017. Output of model simplification, after dropping terms that did not significantly improve the overall model fit using likelihood-ratios.

Initial model:
pH ~ N * Fng * SR * FC + (1|Block) + (1|Combination)

| factor | AIC | LRT | Pr(Chi) |
|---|---|---|---|
| N | -76.77 | 1.978 | 0.16 |
| Fng | -77.41 | 1.333 | 0.248 |
| Fng | -77.41 | 1.333 | 0.248 |
| SR | -76.75 | 0.169 | 0.681 |
| FC | -74.92 | 1.526 | 0.466 |
| Fng:SR | -72.44 | 1.857 | 0.173 |
| N x SR | -72.3 | 0.927 | 0.336 |
| FC x SR | -71.23 | 0.847 | 0.655 |
| N x FC | -68.07 | 0.201 | 0.904 |
| N x FC x SR | -64.27 | 3.347 | 0.188 |
| Fng x FC | -63.62 | 0.073 | 0.964 |
| Fng x FC x SR | -59.7 | 1.888 | 0.389 |
| N x Fng | -57.58 | 0.644 | 0.422 |
| N x Fng x SR | -56.23 | 0.761 | 0.383 |
| N x Fng x FC | -54.99 | 1.468 | 0.48 |
| N x Fng x FC x SR | -52.46 | 4.006 | 0.135 |

**Table S14: Negative soil respiration model.** Effect on multifunctionality, calculated with negative soil respiration, of sown species richness (SR), functional diversity (MPD), community weighted mean specific leaf area (SLA), fungicide (Fng), nitrogen enrichment (N), multifunctionality threshold and their interaction. Output of the linear mixed effects model after model simplification (scaled variables).

Initial model:
multifunctionality ~
Threshold * (N + SLA + Fng + SR + MPD) ^2
- Threshold:SLA:MPD - SLA:MPD + (1|Block) + (1|Combination) + (Threshold |Plot_Nr)

| Factor | Estimate | Std.Error | Significance |
|---|---|---|---|
| (Intercept) | 0.591 | 0.013 | |
| Threshold | -0.159 | 0.003 | <0.001 |
| N | 0.013 | 0.005 | |
| Fng | -0.010 | 0.004 | |
| SR | 0.008 | 0.011 | |
| SLA | 0.033 | 0.009 | |
| MPD | 0.001 | 0.008 | |
| **N x MPD** | -0.009 | 0.004 | 0.067 |
| **Fng x MPD** | -0.007 | 0.004 | 0.077 |
| **SR x SLA** | 0.025 | 0.009 | 0.006 |

Conditional R-squared: 0.88

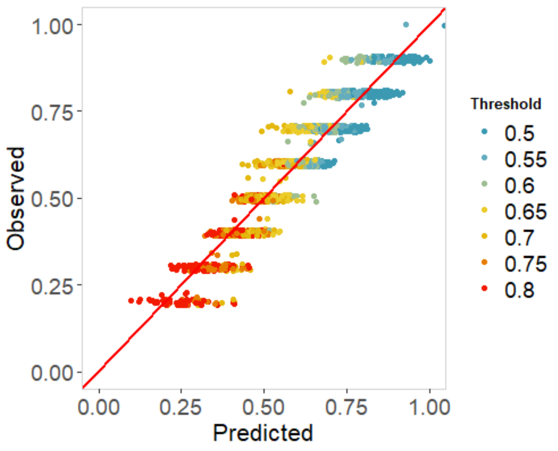


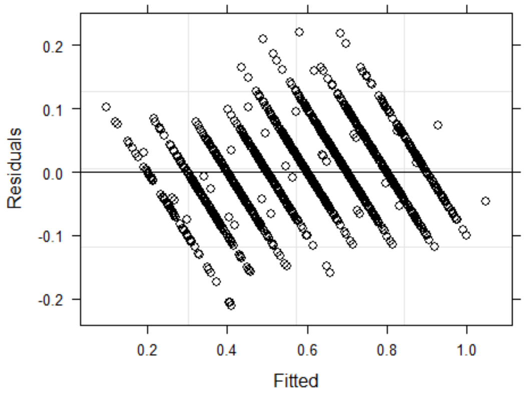


Soil respiration is positively correlated with total plant cover:

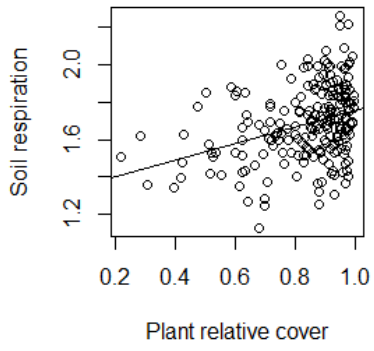

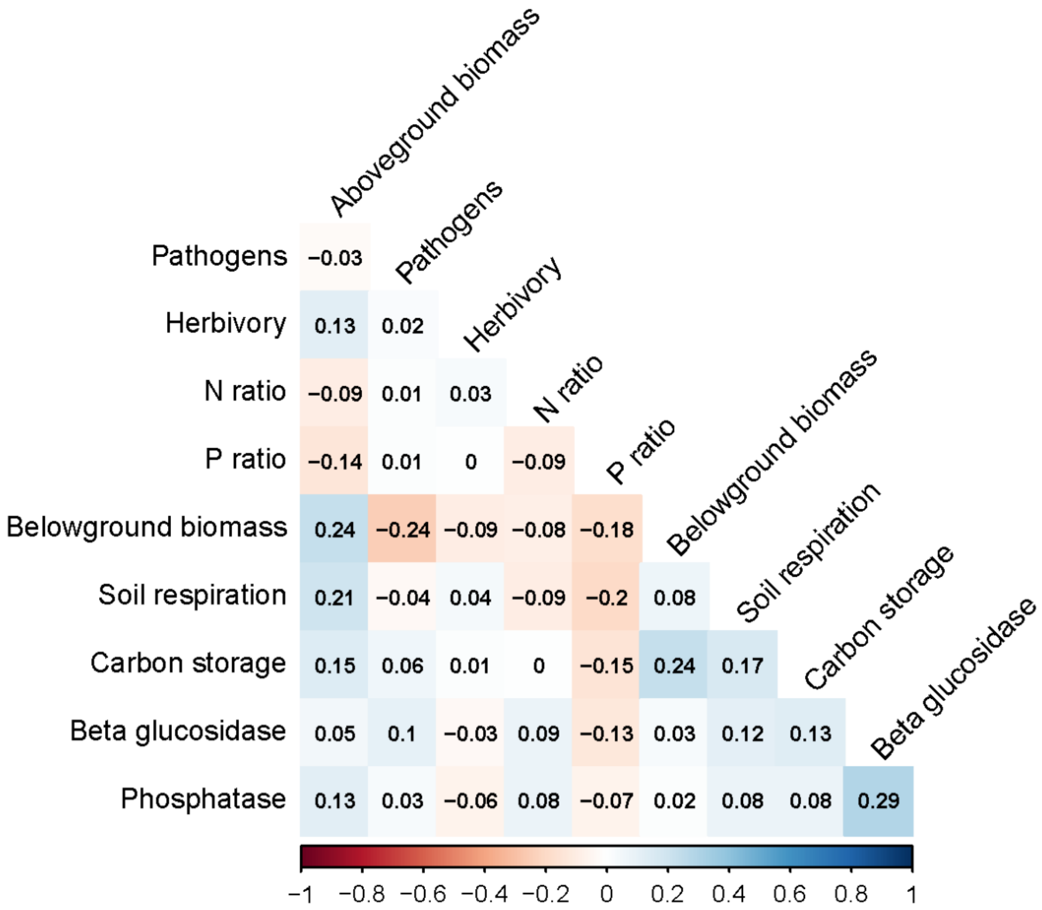


**Figure S8:** Pearson's correlations between the functions measured on 216 plots after log or square root transformation to fit a normal distribution (see Table S9 for the transformation detail). The functions are not strongly correlated with each other, meaning that we did not include redundant functions in our calculation of multifunctionality.

**Table S15: Monoculture/mixture model.** Lmer output correcting for the fact that monocultures are all coded as 0 MPD. The term “mixture” is a categorical variable distinguishing monoculture from mixture (0/1). The mixture term drops out from the final model.

Initial model:
multifunctionality ~
Threshold * (N + SLA + Fng + SR + MPD)^2 - Threshold:SLA:MPD - SLA:MPD
+ mixture:SLA + mixture:N
+ (1|Block) + (1|Combination) + (Threshold |Plot_Nr)

| Factor | Estimate | Std.Error | Significance |
|---|---|---|---|
| (Intercept) | 0.572 | 0.014 | |
| Threshold | -0.179 | 0.004 | <0.001 |
| N | 0.014 | 0.005 | |
| Fng | -0.008 | 0.004 | |
| SR | 0.010 | 0.011 | |
| SLA | 0.033 | 0.009 | |
| MPD | 0.001 | 0.009 | |
| N x MPD | -0.010 | 0.004 | 0.014 |
| Fng x MPD | -0.008 | 0.004 | 0.039 |
| SR x SLA | 0.025 | 0.009 | 0.005 |

Conditional R-squared: 0.88

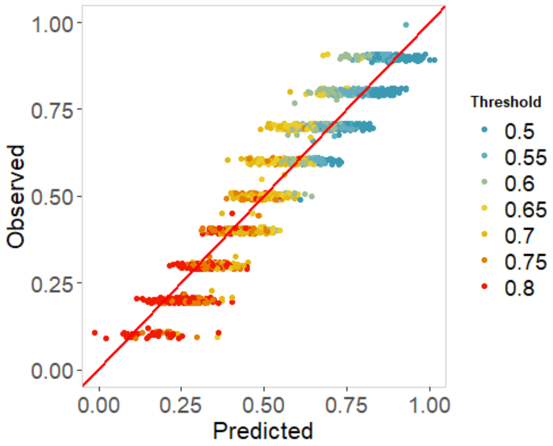


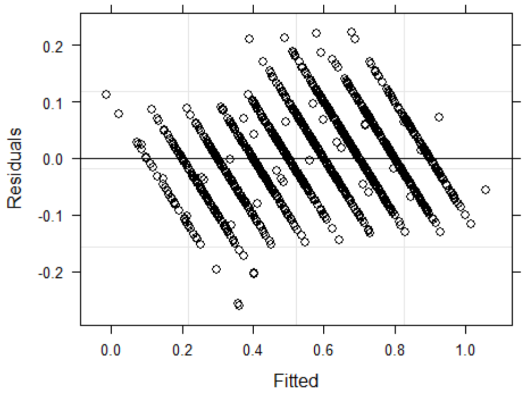
